# JARID2 and AEBP2 regulate PRC2 activity in the presence of H2A ubiquitination or other histone modifications

**DOI:** 10.1101/2020.04.20.049213

**Authors:** Vignesh Kasinath, Curtis Beck, Paul Sauer, Simon Poepsel, Jennifer Kosmatka, Marco Faini, Dan Toso, Ruedi Aebersold, Eva Nogales

**Affiliations:** QB3 Institute, Department of Molecular and Cell Biology, University of California, Berkeley, California 94720, USA; Department of Molecular and Cellular Biology, University of California, Berkeley, California 94720, USA; Howard Hughes Medical Institute, University of California, Berkeley, California 94720, USA; Center for Molecular Medicine Cologne, University of Cologne, Cologne, Germany; Cologne Excellence Cluster for Cellular Stress Responses in Ageing-Associated Diseases (CECAD), University of Cologne, Cologne, Germany; Department of Biology, Institute of Molecular Systems Biology, ETH Zurich, 8093, Switzerland; Faculty of Science, University of Zurich, Zurich, Switzerland; Molecular Biophysics and Integrated Bioimaging Division, Lawrence Berkeley National Laboratory, USA

## Abstract

The Polycomb repressive complexes PRC1 and PRC2 functionally interact to coordinate cell type identity by the epigenetic regulation of gene expression. It has been proposed that PRC2 is recruited to genomic loci via the recognition of PRC1-mediated mono-ubiquitination of histone H2A at lysine 119 (H2AK119ub1), but the mechanism of this process remains poorly understood. Here, we report the cryo-EM structure of human PRC2 with cofactors JARID2 and AEBP2 bound to a nucleosome substrate containing H2AK119ub1. We find that JARID2 and AEBP2 each interact with one of the two ubiquitin molecules in the nucleosome. A ubiquitin-interaction motif (UIM) in JARID2 is sandwiched between ubiquitin and the histone H2A-H2B acidic patch. Simultaneously, the tandem zinc-fingers of AEBP2 interact with the second ubiquitin and the histone H2A-H2B surface on the opposite side of the nucleosome. JARID2 plays a dual role in the H2AK119ub1 dependent stimulation of PRC2 through interactions with both EED via its K116 trimethylation and with the H2AK119-ubiquitin. AEBP2, on the other hand, appears to primarily serve as a scaffold contributing to the interaction between PRC2 and the H2AK119ub1 nucleosome. Our structure also provides a detailed visualization of the EZH2-nucleosome interface, revealing a segment of EZH2 (named “bridge helix”) that is stabilized as it bridges the EZH2(SET) domain, the H3 tail and the nucleosomal DNA. In addition to the role played by AEBP2 and JARID2 in PRC2 regulation by H2AK119ub1 recognition, we also observe that the presence of these cofactors partially overcomes the inhibitory effect that H3K4- and H3K36-trimethylation have on core PRC2. Together, our results reveal the central role played by cofactors JARID2 and AEBP2 in orchestrating the crosstalk between histone post-translational modifications and PRC2 methyltransferase activity.

---

The Polycomb complexes PRC1 and PRC2 are chromatin modifiers critical for lineage commitment during embryonic development and for maintaining cell type identity post-differentiation(*1, 2*). PRC1 is an E3 ubiquitin ligase that catalyzes the mono-ubiquitination of histone H2A at lysine 119 (H2AK119ub1) through its RING1A/B subunit(*3, 4*). The core complex of PRC2 (PRC2-core) contains four proteins (EZH2, EED, SUZ12 and RBAP46/48) and catalyzes the mono-, di- and tri-methylation of histone H3 at lysine 27 (H3K27me1/2/3)(*5*). Accessory subunits, also termed cofactors, associate with the PRC2-core to constitute PRC2 variants with distinct biological functions(*6*). Two mutually exclusive forms of PRC2 have been reported, sometimes referred to as PRC2.2 (containing JARID2 and AEBP2) or PRC2.1 (containing PHF1, MTF2, PHF19 or EPOP/PALI1)(*4, 6*). PRC2 cofactors have been proposed to have roles in both the initial recruitment of the complex to chromatin and the allosteric regulation of its HMTase activity through interactions with long-non coding RNAs (lncRNAs), GC-rich DNA, and the histone mark H2AK119ub1(*7-9*). Recent studies have suggested a highly regulated hierarchical recruitment model involving a complex crosstalk between the activities of the different PRC1 and PRC2 complexes in the establishment of polycomb heterochromatin domains(*3, 10*). The loss of H2AK119ub1 on polycomb target genes has been observed to result in a corresponding reduction in H3K27me3 deposition, suggesting a strong connection between PRC1 and PRC2 catalytic activities(*11, 12*). However, the specific role played by PRC1-mediated H2AK119ub1 in the recruitment of PRC2 has remained elusive, and there have been conflicting observations reporting both inhibition and activation of PRC2 by this histone mark(*13, 14*). H2AK119ub1 nucleosomes are recognized by PRC2 containing AEBP2 and JARID2, which are thought to be involved in both the H2AK119ub1 dependent recruitment and the regulation of HMTase activity(*13*). Recent studies have shown that this recruitment is mediated by a putative ubiquitin interaction motif (UIM) located near the N-terminus of JARID2(*7*). Despite these biochemical insights, the mechanistic details of H2AK119ub1 recognition by PRC2 have remained unknown. In addition to their role in H2AK119ub1 recognition, core PRC2 together with AEBP2 and JARID2 colocalize to CpG rich promoter regions with H3K4me3, as well as bivalent domains containing both H3K4me3 and H3K27me3 in embryonic stem cells(*15-19*). It has been accepted in the field that active transcription marks (H3K4me3 and H3K36me3) inhibit PRC2 HMTase activity; however, the effect of cofactors AEBP2 and JARID2 in the H3K4me3- and H3K36me3-mediated inhibition remains poorly understood(*20, 21*). Using cryo-EM and biochemical assays, we provide mechanistic insight into how the cofactors AEBP2 and JARID2 specifically recognize H2AK119ub1, and how they affect PRC2 HMTase activity in the context of H2AK119ub1 or H3K4/36me3-containing nucleosomes.

We have obtained the cryo-EM structure (3.5 Å) of a 1:1 stoichiometric complex of the complete human PRC2 with cofactors AEBP2 and JARID2 (aa 1-450) (referred to, from here on, as PRC2-AJ_1-450_) bound to a H2AK119ub1-containing nucleosome (Ncl-ub) (Figure 1, fig. S1-4, Movie 1). The overall architecture of the Ncl-ub bound PRC2-AJ_1-450_ agrees very well with our previous cryo-EM structure of PRC2 containing AEBP2 and JARID2(*22*). The engagement of the nucleosome also agrees with what we visualized for the substrate nucleosome in the context of the five-subunit PRC2-AEBP2 complex bound to a dinucleosome(*23*). Improvements in cryo-EM sample preparation that overcame the need for crosslinking and stabilized the structure from the deleterious effect of the air-water interface, resulted in an intact PRC2 bound to the nucleosome, allowing us to structurally model several new regions of the complex (Figure 1, Movie 1). Our study provides the structures of JARID2 and AEBP2 segments recognizing the ubiquitinated nucleosome, of regions of EZH2 linking the SANT2 and the SET domain, a near complete model for SUZ12, and new segments of AEBP2. In addition to this 1:1 stoichiometry, our cryo-EM data processing also identified a small sub-population with two PRC2-AJ_1-450_ each engaging one of the two histone H3 tails, on the opposite sides of the nucleosome (fig. S5). The two PRC2 complexes in that arrangement do not visibly interact with each other and therefore likely represent independent binding events.

**Figure 1.**
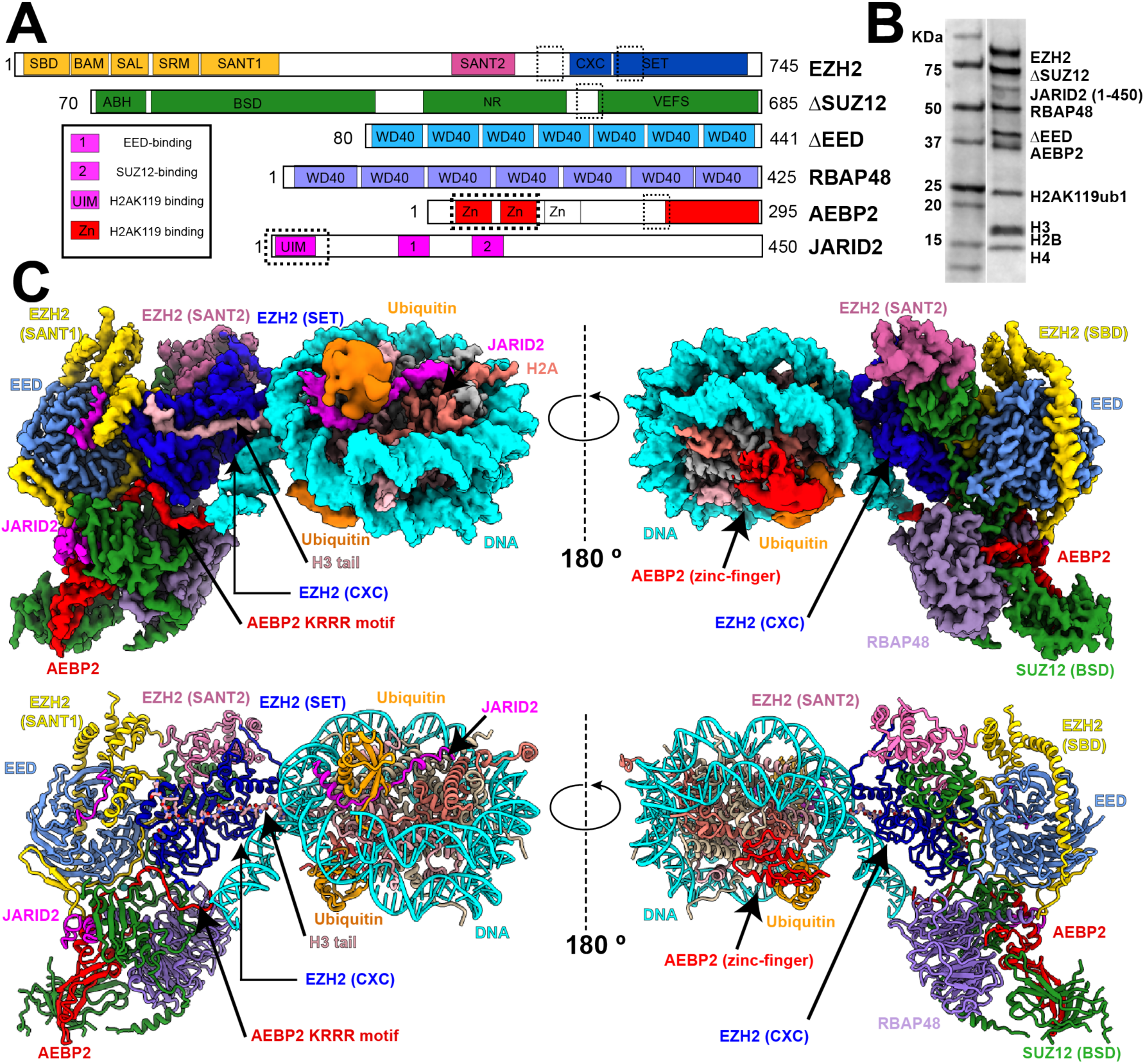
Cryo-EM structure of PRC2-AEBP2-JARID2 bound to a H2AK119ub1 Nucleosome. (A) Schematic representation of the proteins in the PRC2-AEBP2-JARID2 complex. New modeled regions in EZH2, JARID2 and AEBP2 that contribute to the interaction with the nucleosome are marked by dashed boxes. (B) Coomassie stained gel showing all the PRC2 core subunits, cofactors JARID2 and AEBP2, and histone proteins in the sample used for structural studies. (C) (Top) Cryo-EM density map for PRC2-AJ_1-450_ bound to an H2AK119ub1-containing nucleosome. (Bottom) Atomic model of PRC2-AJ_1-450_ bound to an H2AK119ub1-containing nucleosome, with EZH2(SANT1) highlighted in gold, EZH2(SANT2) in hot pink, EZH2(SET) in blue, EED in light blue, RBAP48 in light purple, SUZ12 in green, JARID2 in magenta, AEBP2 in red, ubiquitin in orange, histone H3 in light pink, H2A in salmon, H4 and H2B in khaki and nucleosome DNA in cyan. This color-coding is consistent with our previous structural studies and also applies to Movie 1.

Our structure shows that the SET and CXC domains of EZH2 bind the H2AK119ub1 nucleosome, which contains unmodified histone H3K27, using the same interaction interface previously shown for the substrate nucleosome in our cryo-EM reconstruction of PRC2 bound to a dinucleosome(*23*) (Figure 1C, fig. S6). The nucleosomal DNA is further contacted by the KRRKLKNKRRR motif (aa 171-181) of AEBP2, in agreement with previous studies suggesting its potential role in DNA interaction(*17, 24*) (fig. S6). Cryo-EM analysis of the conformational heterogeneity in the sample revealed that these two PRC2-nucleosome interfaces are flexible (fig. S6). In the present structure we have now been able to identify and model a region of EZH2 rich in lysines and arginines (aa 487-513) that interacts extensively with the nucleosomal DNA. The region includes lysines K509 and K510 within the IQLKK motif of EZH2, which had been predicted to be disordered and shown to be modified during PRC2 auto-methylation(*25, 26*). Mass spectrometry analysis of our PRC2-AJ_1-450_ sample confirms the presence of dimethylation of both K509 and K510. When bound to substrate nucleosome, the aa 497-511 segment in EZH2 forms an alpha helix that is sandwiched between the nucleosomal DNA, the EZH2 (SET) domain, and the H3 tail and that we therefore term the EZH2 bridge helix (Figure 2A). Its tripartite interaction results in the stabilization of this EZH2 region that is otherwise unstructured or highly flexible in previous structural studies of PRC2 lacking nucleosome substrate(*22, 27, 28*). The resolution of the map allowed us to build a large region of the histone H3 tail (aa 21-40) *de novo* and to describe a network of van der Waals and hydrogen bonding interactions that guide the H3 tail into the catalytic site of EZH2 (Figure 2B). Of notice, cancer mutations frequently observed in the EZH2(SET) and EZH2(CXC) domains map to regions that we now see interacting with the histone H3 tail and DNA (Figure 2C). Our findings extend our current knowledge on substrate nucleosome engagement, providing a complete atomic model of DNA and histone tail contacts by EZH2.

**Figure 2.**
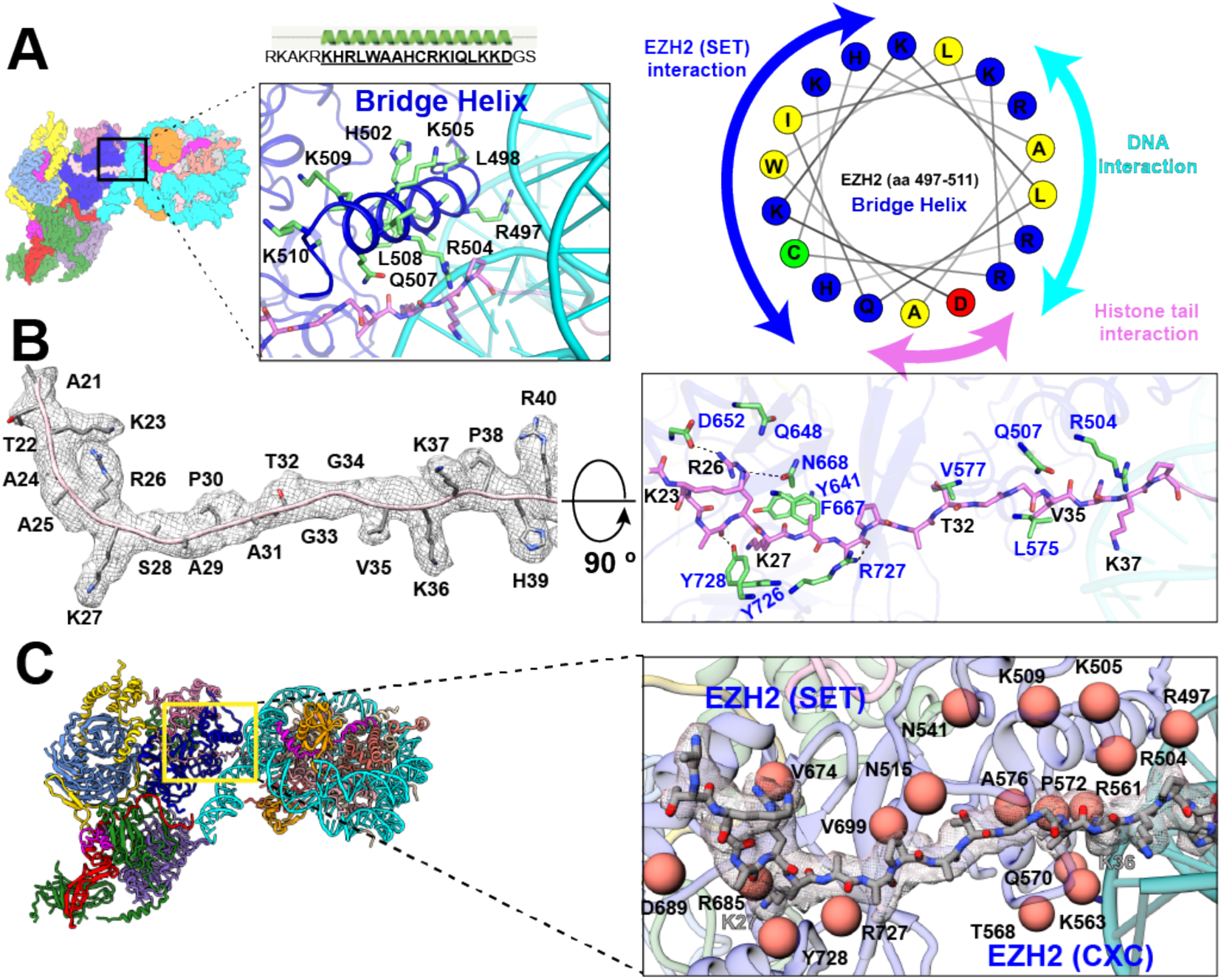
Interaction of EZH2 with the histone H3 tail and nucleosomal DNA. (A) (Top) Cartoon representation of the newly defined EZH2 (aa 497-513) bridge helix that interacts with nucleosomal DNA and the H3 tail. (Left) Cryo-EM structure of the PRC2-AEBP2-JARID2 complex bound to an H2AK119ub1-containing nucleosome. The zoom out shows the bridge helix, with residues interacting with the EZH2 (SET) domain, nucleosomal DNA and the histone tail depicted in stick representation. (Right) Helix wheel diagram for the bridge helix showing the distribution of positively charged residues on the nucleosomal DNA interacting face (cyan), positive and hydrophobic residues on the EZH2 (SET) interacting face (blue), and the residues interacting with the backbone of the H3 tail (pink). (B) (Left) Density map of the histone H3 tail (aa 21-40) with the corresponding atomic model. Clear density for residues R26 and K27, which have been previously observed, as well as density for K23, K36, K37 and V35, allow to define the full extent of interaction between EZH2 (SET) and the histone H3 tail. (Right) Extensive electrostatic and van der Waals interaction between the residues in EZH2 (SET) (blue; shown in green stick representation) and the histone H3 tail (black; shown in pink stick representation) guide the H3 tail into the catalytic site. (C) Close-up view of the interaction of EZH2 (SET, CXC) with the histone H3 tail (stick representation with corresponding cryo-EM density in transparency) showing as orange spheres residues involved in either direct or indirect interaction with the histone H3 tail and nucleosome DNA that are also frequently found mutated in cancers (COSMIC database(*39*)).

Proximal to the EZH2(SET) domain, an N-terminal segment of JARID2 (aa 24-57) is sandwiched between one of the two ubiquitin molecules and the H2A-H2B acidic patch (Figure 3A). The JARID2 segment aa 24-39, which corresponds to the reported UIM, interacts extensively with both the ubiquitin and nucleosomal DNA. The ubiquitin conjugated to H2AK119C is oriented such that the JARID2 UIM interacts with the hydrophobic patch on ubiquitin that includes I44, which had previously been shown to be important for JARID2 interaction with ubiquitin(*7*) (fig. S7). Additionally, the following JARID2 (aa 40-57) segment, containing positively charged arginine and lysine residues, interacts with the H2A-H2B acidic patch (Figure 3A). A cryo-EM reconstruction of PRC2-AEBP2-JARID2 (aa 106-450) (referred to as PRC2-AJ_106-450_) that lacks aa 24-57 of JARID2 also lacks the density we assigned to the JARID2 UIM when bound to Ncl-ub, confirming our model (fig. S8). In the absence of the JARID2 (aa 24-57) segment, the ubiquitin conjugated to H2AK119C is conformationally flexible and therefore becomes invisible in our cryo-EM reconstructions (fig. S8).

**Figure 3.**
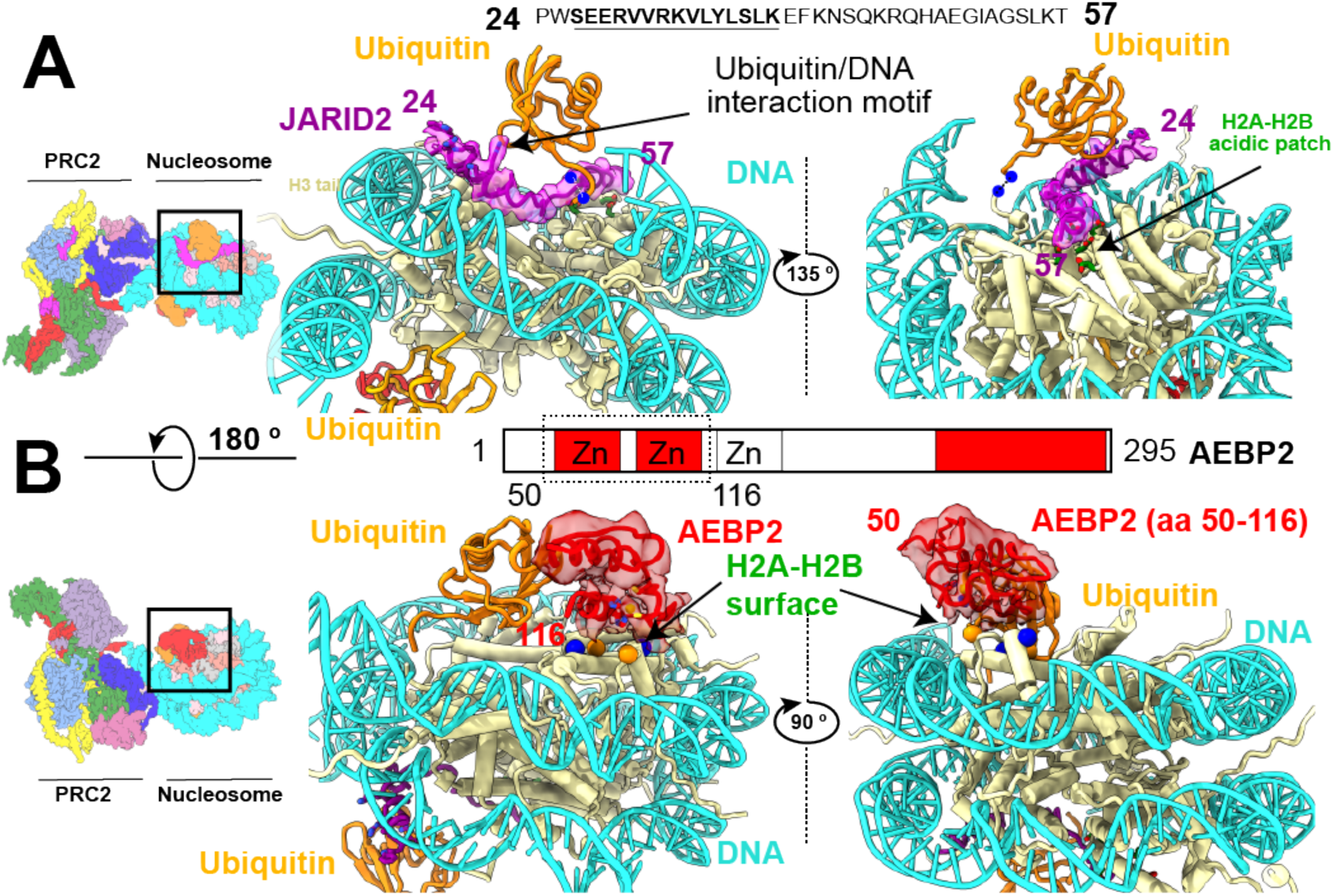
JARID2 and AEBP2 interact with ubiquitin and H2A-H2B surface. (A) Cryo-EM density and model of JARID2 (aa 24-57) (magenta) showing how this segment of JARID2 is sandwiched between ubiquitin and the histone surface. The JARID2 UIM (aa 24-41) forms an alpha helix that interacts with both ubiquitin and nucleosomal DNA. JARID2 (aa 42-57), which contains positively charged arginine and lysine residues, sits on top of the conserved H2A-H2B acidic patch. The blue spheres with dashes lines connecting them indicate the H2AK119C-G76C conjugation. (B) The tandem C2H2 zinc fingers of AEBP2 (aa 50-116) interact with ubiquitin, the H2A-H2B surface, and nucleosomal DNA. The second AEBP2 zinc finger (aa 85-116) interacts extensively with a H2A-H2B surface that containing a mixture of positive and negatively charged residues (blue and orange balls).

On the opposite, distal face of the nucleosome, the first two of the three tandem C2H2 zinc-fingers of AEBP2 (aa 50-116) interact with ubiquitin and with an H2A-H2B surface containing both acidic and basic residues (Figure 3B). The interaction surface between the H2A-H2B and the AEBP2 zinc fingers is similar to that observed for other zinc fingers interacting with the histone surface(*29, 30*) (fig. S7). The cryo-EM reconstruction of PRC2-AJ_1-450_ bound to an unmodified nucleosome (lacking H2AK119ub1) shows no density for either the JARID2 aa 24-57 or the AEBP2 tandem zinc-finger regions, indicating that the interactions of these motifs with the H2A-H2B surface is weak in the absence of ubiquitin (fig. S8). This is further confirmed by cryo-EM reconstructions of PRC2-AEBP2 bound to an unmodified nucleosome (fig. S8). In conclusion, our cryo-EM structures show that both JARID2 and AEBP2 mediate the recognition of H2AK119ub1, providing additional anchoring interactions with the nucleosome besides those of EZH2 with nucleosomal DNA and the H3 tail, and thus supporting a mechanism of JARID2- and AEBP2-dependent recruitment of PRC2 to PRC1-modified nucleosomes.

Given the role of JARID2 and AEBP2 in H2AK119ub1 recognition and recruitment to chromatin, and the conflicting data on the effect of this modification on PRC2 HMTase activity, we decided to compare the enzymatic activity of PRC2 complexes in the presence or absence of these cofactors. We carried out activity assays for four different PRC2 complexes, PRC2-core, PRC2 AEBP2, PRC2-AJ_106-450_, PRC2-AJ_1-450_, on either unmodified mono-nucleosomes or Ncl-ub (Figure 4A, B, fig. S9, 10). We performed our HMTase activity assays probing for both methyltransferase activity based on total SAH production (thus quantifying H3K27me1, H3K27me2 and H3K27me3 cumulatively), and for H3K27me3 specifically using western blot analysis. In agreement with previous results, our analysis of HMTase activity shows that PRC2 containing both AEBP2 and JARID2 is a much more active methyltransferase compared to either PRC2 containing AEBP2 alone or PRC2 core, both of which exhibit poor HMTase activity(*31*) (Figure 4A, B). As expected, the core PRC2 does not distinguish the ubiquitinated from the unmodified nucleosome. Despite the interaction of the AEBP2 tandem zinc-fingers with both the H2AK119ub1 and H2A-H2B surfaces, the HMTase activity of PRC2-AEBP2 on unmodified nucleosomes was only slightly lower (based on total methylation) to that on Ncl-ub. A PRC2-AJ_106-450_, containing the JARID2 K116me3 stimulatory region but lacking the H2AK119ub1 recognizing UIM, displayed enhanced but otherwise similar HMTase activity on either unmodified nucleosomes or H2AK119ub1-containing nucleosomes, as expected based on our structure. The inclusion of the larger JARID2 version (aa 1-450), containing the H2AK119ub1-recognizing UIM in addition to the stimulatory JARID2 K116me3 segment, results in a further stimulation of PRC2 HMTase activity on H2AK119ub1-containing nucleosomes compared to unmodified nucleosomes (based both on total methylation and on specific trimethylation). Taken together, our results suggest that the primary allosteric activation of PRC2 by JARID2 involves the binding of JARID2 K116me3 to EED, with the subsequent stabilization of EZH2(SET) catalytic site, with a secondary, modest contribution from the JARID2 UIM interaction with H2AK119ub1 and the H2A-H2B acidic surface. AEBP2 likely plays a smaller role during the recruitment of PRC2 via H2AK119ub1 recognition. The increase in HMTase activity of PRC2-AJ_1-450_ on Ncl-ub could be due to an increased residence time of PRC2 on the nucleosome in the presence of the additional anchoring interactions of the cofactors with Ncl-ub.

**Figure 4.**
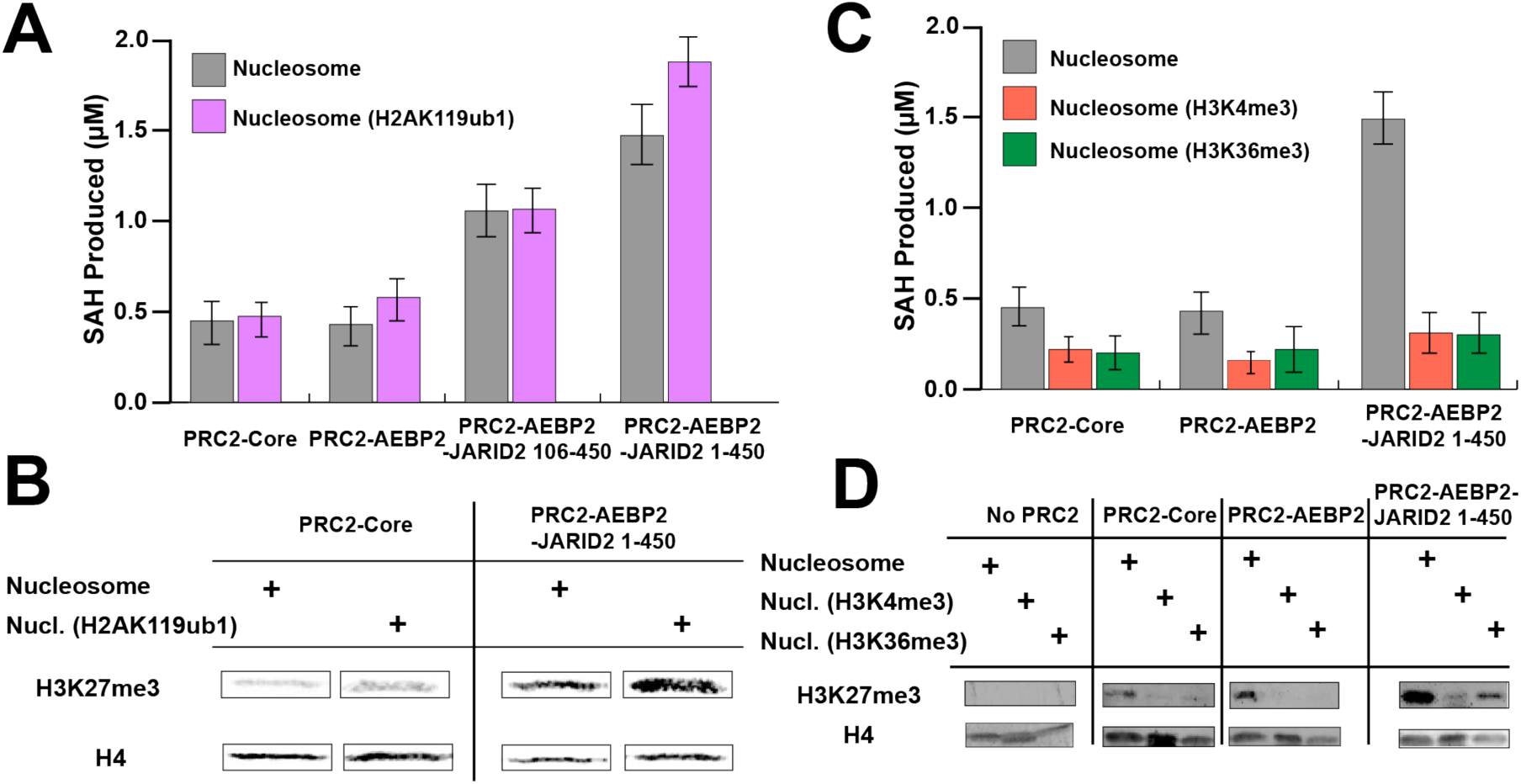
HMTase activity of different PRC2 complexes on modified nucleosomes. (A) Bar graph showing the comparison of end-point, cumulative HMTase activity (H3K27me/me2/me3) of PRC2 core, PRC2-AEBP2, PRC2-AEBP2-JARID2 (106-450), and PRC2-AEBP2-JARID2 (1-450) on unmodified nucleosomes (grey) and on H2AK119ub1 containing nucleosomes (magenta). (B) Western blot analysis comparing the H3K27me3 production for unmodified nucleosomes versus H2AK119ub1 containing nucleosomes. (C) Bar graph showing end point assays for cumulative H3K27me1/me2/me3 activity for different PRC2 complexes on nucleosomes: unmodified (grey), H3K4me3-containing (orange) and H3K36me3-containing (green). (D) Western blot analysis showing the comparison of H3K27me3 produced by the different PRC2 complexes acting on nucleosomes, either unmodified or containing H3K4/K36me3 modifications.

In view of the involvement of cofactors in the crosstalk between H2AK119ub1 and PRC2 HMTase activity, we asked how the presence of JARID2 and/or AEBP2 could also have an effect on HMTase activity in the presence of the active transcription marks H3K4me3 and H3K36me3. To our knowledge, the *in vitro* effect of these modifications on PRC2 HMTase activity has been reported only for the PRC2 core complex lacking cofactors. Again, we performed HMTase activity assays probing for total methyltransferase activity and for H3K27me3 specifically, and we used three different PRC2 complexes containing either AEBP2 and JARID2 (aa 1-450), AEBP2 alone, or just the PRC2-core. We found that in both types of assays, the HMTase activities of PRC2-core and PRC2-AEBP2 complexes are inhibited by nucleosomes containing either H3K4me3 or H3K36me3, in agreement with previous studies(*20*) (Figure 4C, D, fig. S9, 10). PRC2 complexes containing both AEBP2 and JARID2 (1-450) displayed overall inhibition for cumulative activity (H3K27me1/me2/me3) on nucleosomes containing H3K4me3 or H3K36me3, but still retained partial activity on both based on cumulative methylation, and specially for H3K36me3 based on our H3K27me3 western blot analysis. HMTase activity assays with human nucleosomes containing H3K4me3 or H3K36me3 generated via native ligation mirror the observations for Xenopus nucleosomes containing H3K4me3 or H3K36me3 generated as a methyl-lysine analog(*32*) (fig. S10). Thus, our data shows that in the presence of both AEBP2 and JARID2, there is still a strong inhibitory effect of the two marks but that the activation via JARID2 still results in activity that is increased with respect to that without JARID2. We propose that total PRC2 inhibition at actively transcribing regions may require of a combination of H3K4/36me3- and RNA-mediated inhibition, or perhaps the eviction of PRC2 containing JARID2 and/or AEBP2(*9, 20, 33-35*). To gain some mechanistic understanding of the inhibition observed for H3K36me3-containing nucleosomes, we carried out structural modeling of the H3K36me3 within our cryo-EM structure of PRC2-AJ_1-450_ bound to a H2AK119ub1-containing nucleosome. Modeling suggests that while there is limited space for the bulky trimethyl group, it could still be accommodated via alternative side chain conformations (Figure 5). In addition to possible steric effects of accommodating H3K36me3, the loss of electrostatic interaction between the H3K36 and the phosphate backbone of nucleosomal DNA may also contribute to the inhibition of HMTase activity by H3K36me3.

**Figure 5.**
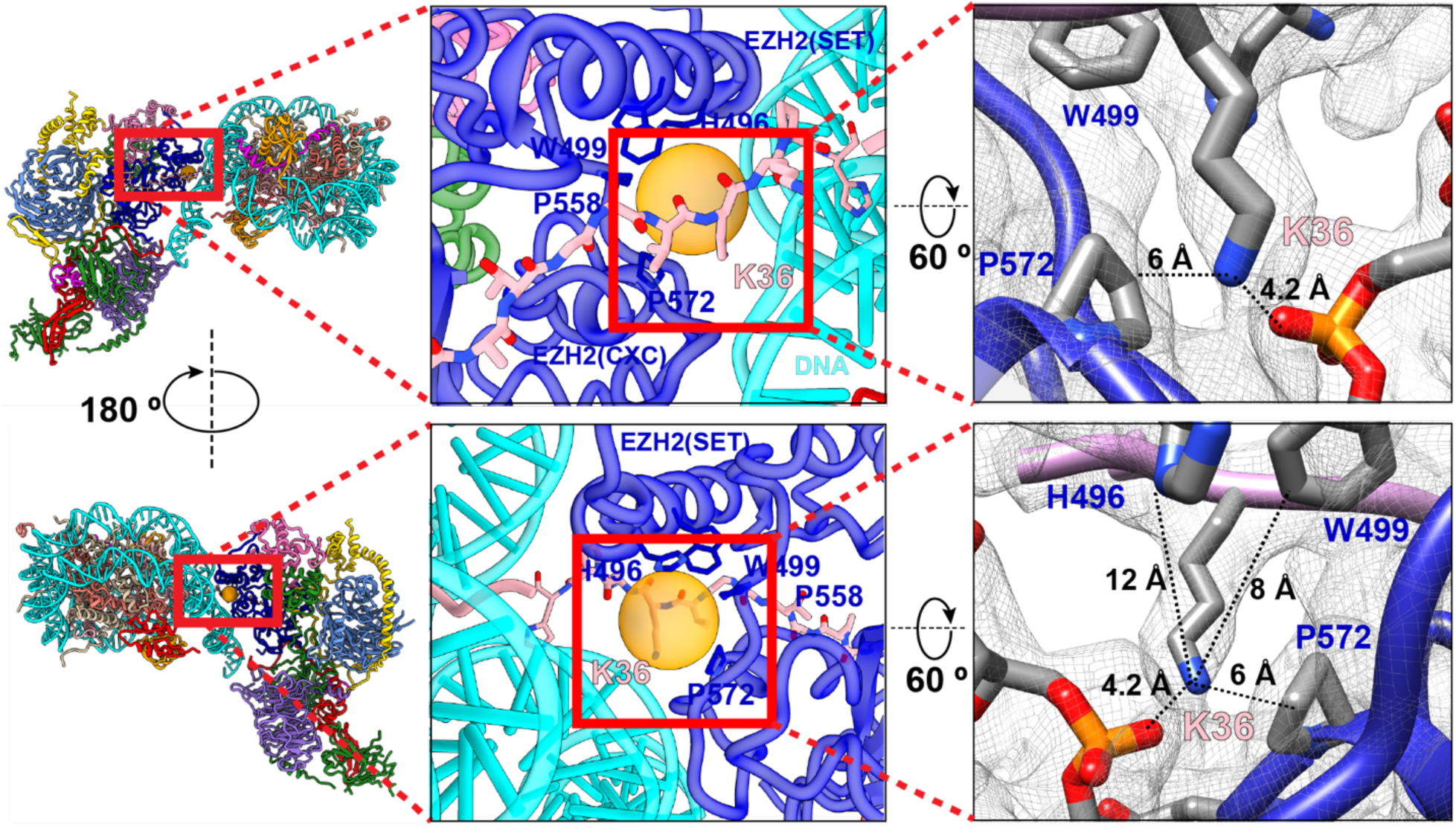
Location of H3K36 at the PRC2-nucleosomal DNA interface. Close-up view of the location of histone H3K36 and its interactions with nucleosomal DNA and PRC2. Bulky residues in PRC2 close to H3K36 are shown in stick representation, with the orange sphere indicating the potential space available for alternative conformations of the H3K36 side chain. On the right, cryo-EM density with the atomic model, showing the distances between H3K36 terminal amine group and nearest bulky residues.

We have obtained a cryo-EM reconstruction (8 Å) of PRC2-AJ_1-450_ bound to a human H3K4me3-containing nucleosome (fig. S11, 12). Our cryo-EM reconstruction shows that the engagement of PRC2 with this H3K4me3-modified nucleosome is similar to that seen with the H2AK119ub1-containing nucleosome (fig. S12). However, we do not observe any density for the histone H3 tail engaging with the catalytic site, while we notice subtle differences near EED, where the allosteric activator segments (JARID2 K116me3 or H3K27me3) bind (fig. S11, 12). We hypothesize that a possible disruption of the allosteric activation via EED, together with increased conformational flexibility of the H3 tail in the presence of H3K4me3 (as observed in other histone post-translational modifications) could contribute to the H3K4me3-mediated inhibition of PRC2 HMTase activity(*36*). This effect could be more pronounced for PRC2-core due to its increased conformational flexibility in the absence of cofactors, in agreement with previous studies(*20, 37*). Since PRC2 is always associated with different cofactors *in vivo*,, more detailed structural and biochemical analyses of PRC2 in complex with cofactors and bound to H3K4me3 or H3K36me3-containing nucleosomes will be necessary to better understand the inhibitory effect of these marks on PRC2 activity and the partial amelioration of that inhibition by cofactors.

In summary, our structural work has identified a critical region, the EZH2 bridge helix, for the binding of PRC2 to nucleosomes and its engagement of the H3 tail. It also defines the details of how PRC2 cofactors JARID2 and AEBP2 recognize nucleosomes containing H2AK119ub1 and thus provides a mechanistic explanation for the recruitment of PRC2 to PRC1-modified genomic regions. Our activity assays indicate that H2AK119ub1 results in a modest increase in HMTase activity in the presence of both cofactors and that their presence slightly reduces the inhibitory effect of H3K4me3 or H3K36me3. We speculate that the reduced inhibition in the presence of cofactors could be important in the context of establishing and maintaining bivalent domains containing both H3K4me3 and H3K27me3 marks in embryonic stem cells and during imprinting(*38*). Taken together, our results emphasize the extent to which cofactors like AEBP2 and JARID2 can serve as critical regulators of PRC2 activity in different chromatin environments. Additional studies will be required to shed further light onto the complex regulatory mechanisms of PRC2 activity.

## ACKNOWLEDGEMENTS

We thank A. Chintangal and P. Tobias for computational support, D. Reif for his contributions during the early stages of the project, R.M. Glaeser and B.G. Han for valuable advice concerning streptavidin grid preparation, Basil Greber for advice during model building, and Jürg Müller and his lab members for insightful discussions.

## Author Contributions

V.K. established experimental procedures, expressed and purified the different complexes, nucleosomes, prepared the samples and cryo-grids, collected, processed the EM data, carried out model building and refinement and performed the activity assays. C.B. and J.K. assisted with the molecular cloning, protein purification, nucleosome production, negative-stain EM sample screening. C.B. assisted V.K. in performing activity assays and making cryo-grids. S.P. initiated the use of streptavidin monolayer to study PRC2-AEBP2 complexes by cryo-EM. S.P. and P.S. provided Xenopus modified nucleosome samples and valuable feedback during the course of the work. S.P and P.S contributed the cryo-EM map of PRC2-AEBP2 bound to an unmodified nucleosome. M.F collected the mass spectrometry data. V. K and M.F analyzed the mass spectrometry data. R.A supervised the mass spectrometry work. V.K. and E.N. conceived the project and analyzed all the data. E.N. supervised the work. V.K. and E.N. wrote the manuscript with input from all the authors.

## Data and material availability

Cryo-EM density maps and fitted models have been deposited in the Electron Microscopy Data Bank (EMDB) and the PDB for the PRC2-AEBP2-JARID2 (1-450) bound to Ncl-ub (EMDB: 21707 and PDB: 6WKR).

## METHODS

### Protein Purification

PRC2 constructs containing full-length EZH2, ΔEED (aa 80-441), ΔSUZ12 (aa 80-685), full length RBAP48, embryonic isoform of AEBP2, and different JARID2 constructs were expressed and purified as previously described(*22*). PRC2 core components together with cofactors strep-tagged AEBP2 and FLAG-tagged JARID2 were assembled into a multibac construct for HighFive insect cell MacroBac baculovirus expression system(*40*) The cells were transfected with baculovirus at 28 °C for 66 hours and frozen in liquid nitrogen until use. PRC2 constructs were purified by resuspending cell pellets in 25 mM HEPES pH 7.9, 250 mM NaCl, 2 mM MgCl_2_, 1 mM TCEP, 0.1% NP40, 10% glycerol supplemented with 10 µM leupeptin, 0.2 mM PMSF, DNAse, and protease inhibitor cocktail (Roche). All purification steps were performed at 4 °C. Post cell lysis, debris were removed by centrifugation at 15,000 RPM for 35 min. The supernatant was incubated with Step-tactin superflow plus resin (Qiagen) overnight (12 hrs) and resin was washed with low salt buffer A (25 mM HEPES pH 7.09, 250 mM NaCl, 2 mM MgCl_2_, 0.01% NP40, 1 mM TCEP, 10% glycerol, 10 µM leupeptin, 0.2 mM PMSF and protease inhibitor cocktail). The resin was further washed with high salt buffer (buffer A with 1 M NaCl). Subsequently, the resin was washed with buffer A without NP40 and eluted with 10 mM Desthiobiotin. The elutes were pooled and incubated with anti-Flag M2 agarose affinity gel and washed with buffer A with 1 M NaCl. PRC2 complexes were eluted in buffer A (no NP40) with 150 µg/ml 3XFLAG peptide and subjected to overnight TEV cleavage to remove all affinity tags. The complexes were further purified over Superose 6 increase (GE healthcare) equilibrated with 25 mM HEPES pH 7.9, 150 mM NaCl, 2 mM MgCl_2_, 10% glycerol and 1 mM TCEP. Protein was flash frozen in liquid nitrogen as single-use aliquots and stored at -80 °C.

### Nucleosome Reconstitution

Xenopus histones (H2A, H2B, H3 and H4) were expressed and purified as described previously(*41*). All the nucleosomal DNA used in this study contain a CpG Island sequence and a 5’ biotin tag and was assembled by large scale PCR and purified over an anion exchange column, and further purified by ethanol precipitation. The 227 bp nucleosome DNA sequence used for all the studies with Xenopus nucleosomes was: 5’(biotin)CACGCGACTGTGTGCCCGTCAGACGC<u>gtgccgaggccgctcaattggtcgtagacagctctagcac cgcttaaacgcacgtacgcgctgtcccccgcgttttaaccgccaaggggattactccctagtctccaggcacgtgtcagatatatacatcctg</u> tatgcatgcatatcattcgatcggagctcccgatcgatgc - 3’. The 601 positioning sequence is underlined and the CG rich sequence used is indicated in uppercase. For unmodified nucleosomes, equimolar amounts of all histones were dialyzed and the octamer was purified over Superdex 200 (GE healthcare). The DNA and octamer was mixed in a 1:1.1 ratio and purified over a BioRad prep cell after overnight gradient salt dialysis, as described previously(*41*). For modified nucleosomes, MLA analogs of H3K4me3 and H3K36me3 were first generated prior to octamer reconstitution and nucleosome generation(*32*). For the generation of ubiquitinated nucleosomes, first equimolar amounts of H2A (K119C) were crosslinked with 6xHis-ubiquitin (G76C) using dichloroacetone and purified under denaturing conditions over a Nickel column, as described previously(*42*). The purified H2A-ub was dialyzed into water and lyophilized for storage. For H2A-ub containing octamer reconstitution, roughly equimolar amounts of H2A-ub were mixed with H2B under denaturing conditions, and the H2Aub-H2B dimer was purified similarly to the octamer as described above. The H3-H4 tetramer was reconstituted and the purified H3-H4 tetramer, H2A-ub-H2B dimer and nucleosomal DNA were mixed in 1.2:2.4:11 ratio and purified using a BioRad Prep cell after overnight salt dialysis. Nucleosomes were stored on ice for up to a month. All human nucleosomes were purchased from Epicypher with the 5’ biotinylated 187 bp DNA sequence containing the 601 positioning sequence: 5’(BioTEG)GGACCCTATACGCGGCCGCCC<u>tggagaatcccggtctgcaggccgctcaattggtcgtagacagctc tacgtggcgaatttgcgtgcatgcgcctgtcccccgcgttttaaccgccaaggggattactccctagtctccaggcacgtgtcagatatatac atcctgt</u>GCCGGTCGCGAACAGCGACC 3’.

### Kinetic Assays

For PRC2 activity assays on different modified mono-nucleosomes roughly two-fold dilutions of nucleosomes at concentrations from 1 µM to 40 nM were used, together with 20 µM S-adenosyl methionine (SAM). The reaction, carried out at room temperature (RT), was initiated by the addition of PRC2 complexes to a final concentration of 40 nM. 10 µL of the reaction taken at 0.5 min, 1 min and 2 min time points and quenched immediately by addition of trifluoroacetic acid at a final concentration of 0.1 %. The quenched reactions were then transferred to a 384-well white round bottom microplate to profile for methyltransferase activity (cumulative mono-, di- and tri-methylation) using the MTase-Glo assay (Promega), which detects the production of S-adenosyl homocysteine (SAH) as a luminescent indicator. The raw luminescence values were correlated to SAH concentration using standard curves after accounting for basal luminescence of PRC2, nucleosome, and buffer without SAM. The initial rates were determined by linear regression and kinetic parameters determined by fitting the initial rates. All reactions were performed in duplicate and the entire experiment was repeated twice for each PRC2-nucleosome combination. Concentrations lower than 40 nM of PRC2 complex could not be used due to PRC2 dissociation (based on negative stain EM analysis).

Endpoint methyltransferase measurements were performed under similar concentrations, with a 10 µl reaction containing 40 nM PRC2, 1 µM nucleosome, and 40 µM SAM, incubated for 2 hours at RT. The reactions were quenched by addition of trifluoroacetic acid to a final concentration of 0.1%. This was used for luminescent detection using MTase-Glo assay (Promega) as above. Each reaction was done in triplicate and the whole experiment was repeated twice.

### Western Blots

200 nM PRC2 complex was incubated together with 400 nM nucleosomes and 40 µM S-adenosyl methionine in HMTase buffer containing 25 mM HEPES pH 7.9, 50 mM NaCl, 2 mM MgCl_2_, 0.5 mM EDTA, 1 mM DTT at RT for 90 min. The reactions were then run on a 4-20% SDS-Page gel for 30 min at 150 V. Amersham Hybond Nitrocellulose membrane was used for the transfer. Transfers were performed in a 4 °C cold room in an ice bath at 90 V for 10 min followed by 60 V for 30 min. Primary antibodies from Cell Signaling Technology for H3K27me3 (rabbit monoclonal) and H4 (mouse monoclonal) were used. Secondary antibodies from ECL Plex containing Cy5 (goat-anti-rabbit) or Cy3 (goat-anti-mouse) were used to detect H3K27me3 and H4, respectively. This Cell Signaling antibody was highly specific for H3K27me3 on either Xenopus H3 or Human H3.2 or H3.3 containing H3K27me3, with no cross reactivity detected towards H3K4me3 or H3K36me3.

### Cryo-EM sample preparation

Streptavidin monolayer affinity grids were used for cryo-EM grid preparation. Quantifoil Au 2/2 Streptavidin grids were made in-house using procedures described previously(*43*). PRC2-nucleosome complexes were assembled by incubating 200 nM nucleosome with 600 nM PRC2 complex and 80 µM SAH in 25 mM HEPES pH 7.9, 40 mM KCl, 1 mM TCEP, 1 mM MgCl_2_ for 30 min at RT. 3 µl of the complex was then incubated with streptavidin monolayer affinity grids (prewashed in the buffer above) and incubated for 3-8 min in a humidified chamber. The grid was washed twice with 10 µl wash buffer containing 25 mM HEPES pH 7.9, 40 mM KCl, 1 mM MgCl_2_, 1 mM TCEP, 4 % Trehalose and 0.01 % NP40 or 0.05 % b-OG. Following the washes, the buffer was wicked away using Whatman filter paper and 2 µl of wash buffer above was added immediately. The grid was then transferred to the FEI Mark IV Vitrobot and manually blotted for 2 sec at 5 °C and 100 % humidity and plunged into liquid ethane.

### Cryo data collection and processing

All the data sets were collected in Super-Resolution Counting mode with image shift and active beam-tilt correction. All but one data set was collected on either on a Titan Krios or Talos Arctica equipped with Gatan K3 direct detector. One of the data set was collected on a Titan Krios equipped with a Gatan K2 detector. All data were acquired as dark subtracted non-gain corrected movies and gain correction was applied during motion correction using MotionCor2(*44*). Data acquisition was performed using SerialEM using custom macros for automated data collection. A GIF quantum energy filter was used for collection with a 20 eV slit width. The total dose for all the data sets was 50 e^-^/Å^2^. All other parameters for data collection can be found in Table S1.

Negative stain on the PRC2-nucleosome complexes was performed by applying 3-4 µl of 50 nM nucleosome plus 150 nM PRC2 on glow discharged 400 mesh C-flat continuous carbon grids (Protochips) and incubated for less than 20 seconds before staining. Negative stain screening to gauge the intactness of PRC2 was performed by diluting the PRC2 complexes to the appropriate concentration and immediately applying 3-4 µl to the glow discharged grids. All negative stain samples were visualized on a Tecnai F20 microscope operated at 120 kV, using a Gatan Ultrascan 4000 camera, at a nominal magnification of 80000x (1.5 Å/pixel), using a dose of 20 e^-^/Å^2^.

All data processing was done within RELION(*45*). The movie frames were aligned using MotionCor2(*44*) and CTF parameters fit using GCTF(*43*). The summed aligned movies were used to substract the background streptavidin lattice using in-house scripts. These subtracted summed aligned movies were used in RELION version 3.0 and 3.1 for processing. Several algorithms were tried for particle picking including logpicker, RELION autopick based on 2D templates obtained from a combination of logpicker/manual picking, EMAN neural net picking, and Topaz neural net picking(*46, 47*). Topaz was eventually used as it resulted in the cleanest picks and more importantly particles with different orientations. To pick particles within Topaz, about 200-500 unambiguous particles were carefully picked manually and used to train a neural net model. This neural net model was subsequently applied to the entire data set. Topaz neural net models were generated for each data set individually. Multibody refinement and particle subtraction were performed using the latest build of RELION 3.1. For all data sets, initial models were generated within RELION and used as reference for the first round of 3D classification. Subsequent processing steps used good classes from the initial 3D classification as a reference. Following the initial 3D classification, 2D classification was performed for all the data sets to clean up the particle stack. Subsequently the good particles were subjected to refinement and further 3D classification followed by downstream processing. For the PRC2 bound to Ncl-ub the final set of good particles from both data sets, one collected with a Gatan K3 detector and the other collected with a Gatan K2 detector, was combined and refined, followed by 3D classification and subsequent final MultiBody refinement in RELION. This final combined map was used for model building.

### Model Building

All models were built using the cryo-EM maps of PRC2-AJ_1-450_ bound to Ncl-ub. The coordinates of PRC2-AEBP2-JARID2 (106-450) were used as a starting model from which all the coordinates were adjusted and rebuilt in the new map using COOT(*22, 48*). New regions corresponding to EZH2, AEBP2, SUZ12 were built *de novo* into the EM density in COOT(*48*). Regions of JARID2 assigned after comparison with cryo-EM maps of PRC2-AJ_1-450_ bound to unmodified nucleosomes and PRC2-AEBP2 bound to H2AK119ub1 ubiquitinated nucleosomes were built *de novo* starting from an idealized α-helix. The tandem zinc fingers of AEBP2 were built by first docking each zinc finger structure individually into the cryo-EM density and then adjusting it in COOT(*48, 49*).

The nucleosome was built by first docking the atomic model with the 601 positioning sequence into the EM density and all the histones and DNA were manually adjusted and extended in COOT(*48, 50*). The histone H3 tail reaching into the EZH2 catalytic site was built *de novo* in COOT(*48*). The density of histone H2A reaching into the ubiquitin density was used as a marker for rigid body docking of the structure of ubiquitin, such that G76C of ubiquitin would be ∼ 3 Å from H2AK119C. Similarly for the ubiquitin close to the AEBP2 density on the nucleosome, the density of the H2A tail and the density of the C-terminal tail of ubiquitin were used as markers to rigid body dock the ubiquitin(*29, 51*).

The model for the second PRC2 bound to Ncl-ub was obtained by rigid-body docking the model for PRC2 obtained in this study, and the histone H3 tail was modeled as a Poly-Ala chain in the catalytic site of the second PRC2.

The model for the monomeric PRC2-Ncl-ub structure was subjected to global refinement and minimization in real space using PHENIX(*52*). These were then subjected to manual inspection and adjustment in COOT(*48*) followed by refinement again in PHENIX(*52*). Model overfitting was also evaluated in PHENIX.

### Mass spectrometry

Sample preparation was performed as described previously(*22*). Briefly, 50 µg of each PRC2 complex was cross-linked with either 0.5 mM final concentration of isotope-labelled di-Succinimidyl Suberate (DSS-d_0_, DSS-d_12_) (Creative Molecules Inc.) in a final volume of 37 µl of 25 mM HEPES pH 7.9, 150 mM NaCl, 1 mM TCEP, 5% glycerol, incubated at 37° C for 30 minutes at 500 rpm on a Thermomixer (Eppendorf). All reactions were quenched with 50 mM final concentration of ammonium bicarbonate (NH_4_HCO_3_) and evaporated to dryness in a vacuum centrifuge. The dried pellets were dissolved in 50 µl of 8 M urea, reduced with 2.5 mM TCEP for 30 minutes at 37° C and alkylated with 5 mM iodoacetamide (Sigma-Aldrich) for 30 minutes at room temperature, in the dark. Digestion was carried out after diluting urea to 5 M with 50 mM NH_4_HCO_3_ and adding 1% (w/w) LysC protease (Wako Chemicals) for 2 hours at 37° C and subsequently diluting to 1 M urea with 50 mM NH_4_HCO_3_ and finally adding 2% (w/w) trypsin (Promega) for 14 hours at 37° C. Protein digestion was stopped by acidification with 1% (v/v) formic acid. Digested peptides were purified using Sep-Pak C18 cartridges (Waters) according to the manufacturer’s protocol. Cross-linked peptides were enriched by peptide size-exclusion chromatography (SEC) as previously described(*53*). SEC fractions were then reconstituted in 5% acetonitrile and 0.1% formic acid and analysed in duplicates on an HPLC (Thermo Easy-nLC 1200) coupled to a mass spectrometer (Thermo Orbitrap Fusion Lumos). Analytes were separated on an Acclaim PepMap RSLC column (25 cm x 75 µm, 2 µm particle size, Thermo Scientific) over a 60-minute gradient from 7% to 35% acetonitrile at a flow rate of 300 nl/min. The mass spectrometer was operated in data-dependent acquisition (DDA) mode with MS acquisition in the Orbitrap analyzer at 120,000 resolution and MS/MS acquisition in the linear ion trap at normal resolution after collision-induced dissociation. DDA was set up to select precursors from an MS1 full scan with a charge state of +3 or higher within a 3 sec cycle between MS1 scans and a dynamic exclusion of 30 s.

Raw files were loaded in MaxQuant (v1.5.2.8)(*54*) and searched against a Swiss-Prot canonical human proteome database (including common contaminant proteins), with precursor and fragment tolerance of 20 ppm and 0.5 Da, respectively. LFQ mode was activated. The search was performed in target-decoy mode, including reverse decoys and including carbamidomethylation (C) as fixed modification and the following variable modifications: mono-, dimethyl (KR), tri-methyl (K), oxidation (M) and phosphorylation (STY). Results were filtered at 1% peptide and protein level false discovery rate. Peptide intensities for EZH2 peptides were corrected by the abundance of EZH2 protein in each condition with a custom R script.

## Supplementary Materials for

**Fig. S1.**
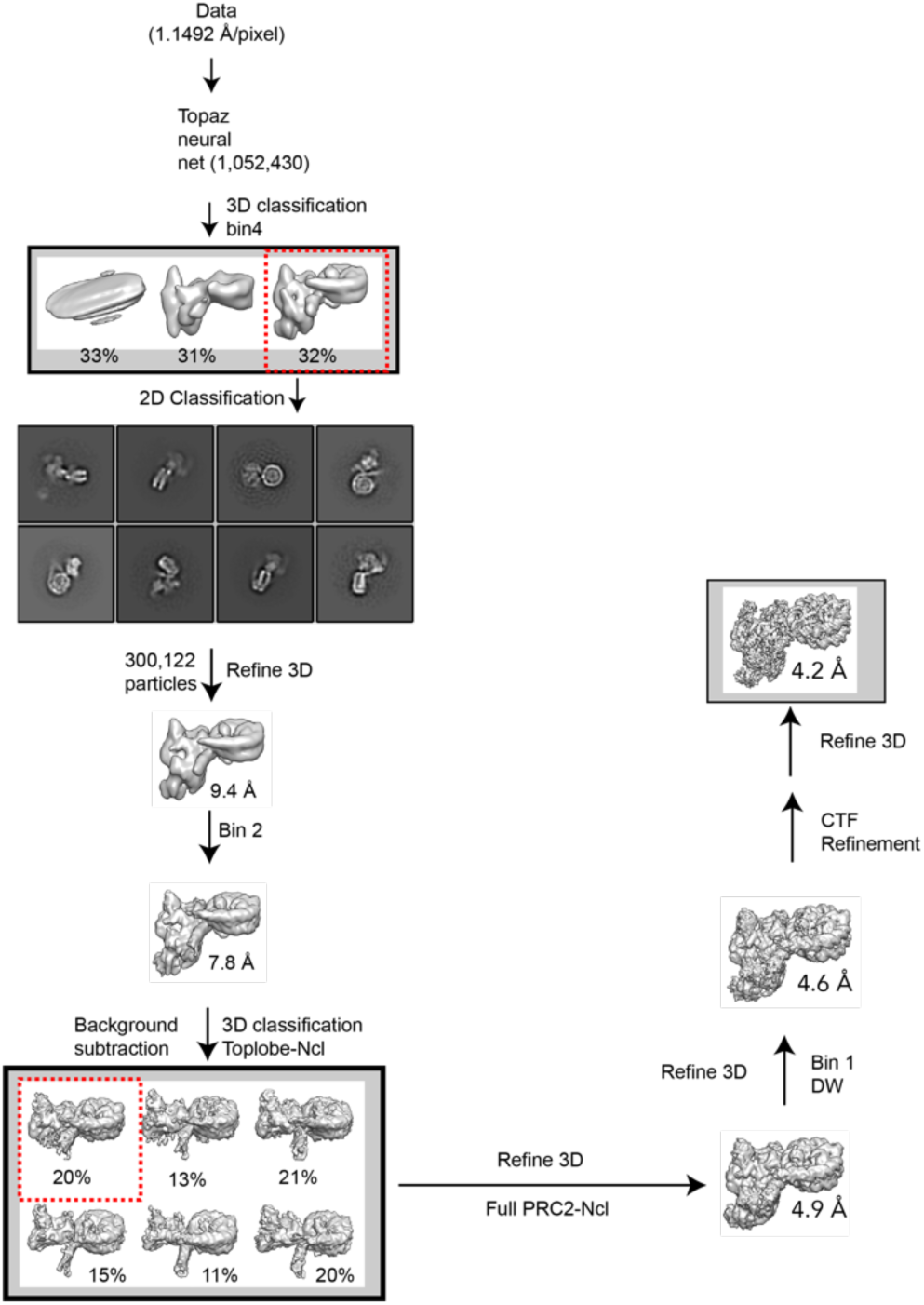
Processing work flow for PRC2-AEBP2-JARID2 (1-450) bound to H2AK119ub containing mono-nucleosome collected on a Titan Krios with a Gatan K2 detector.

**Fig. S2.**
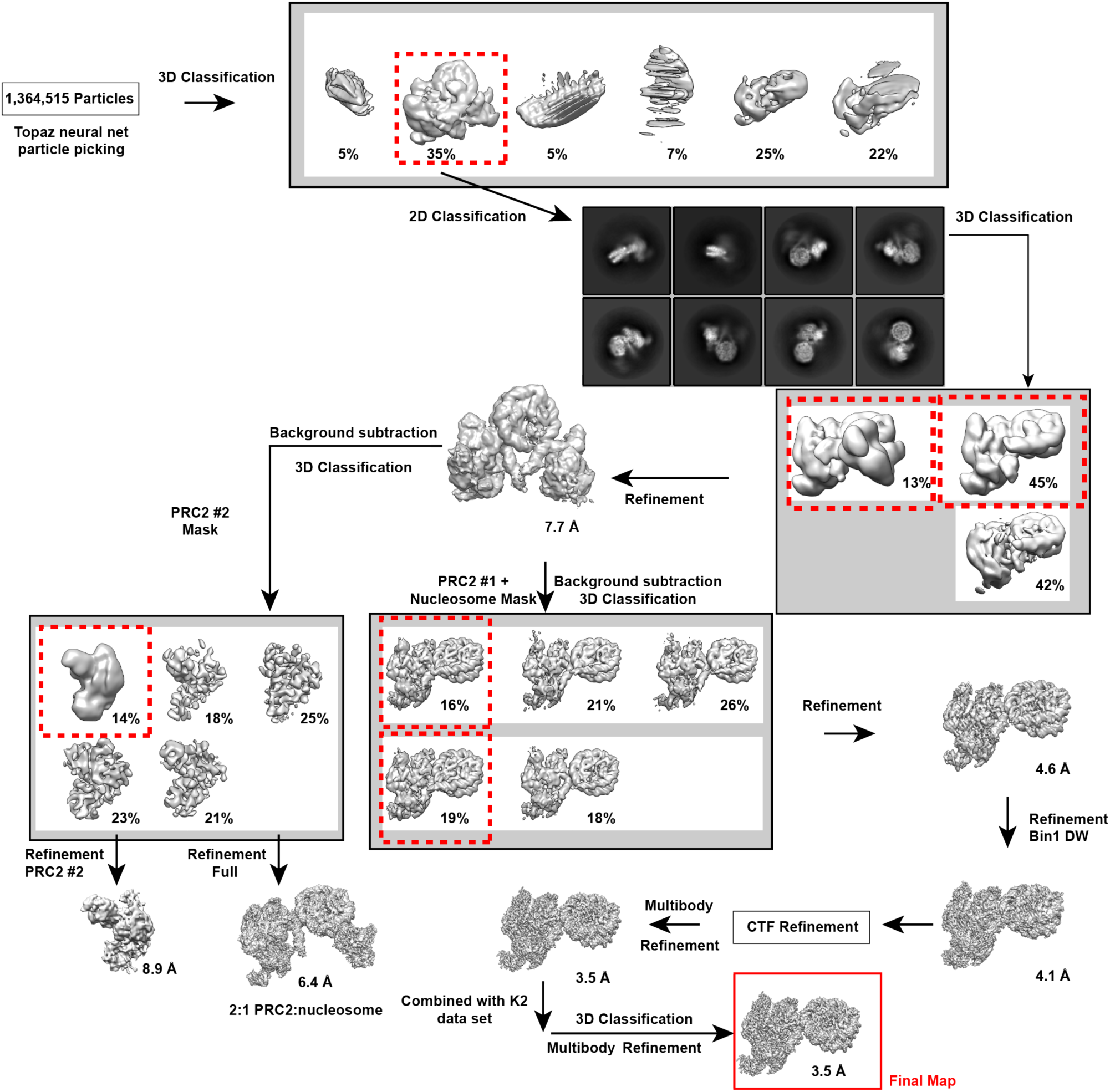
Processing work flow for PRC2-AEBP2-JARID2 (aa 1-450) bound to H2AK119ub containing mono-nucleosome collected on a Titan Krios with Gatan K3 camera.

**Fig. 3.**
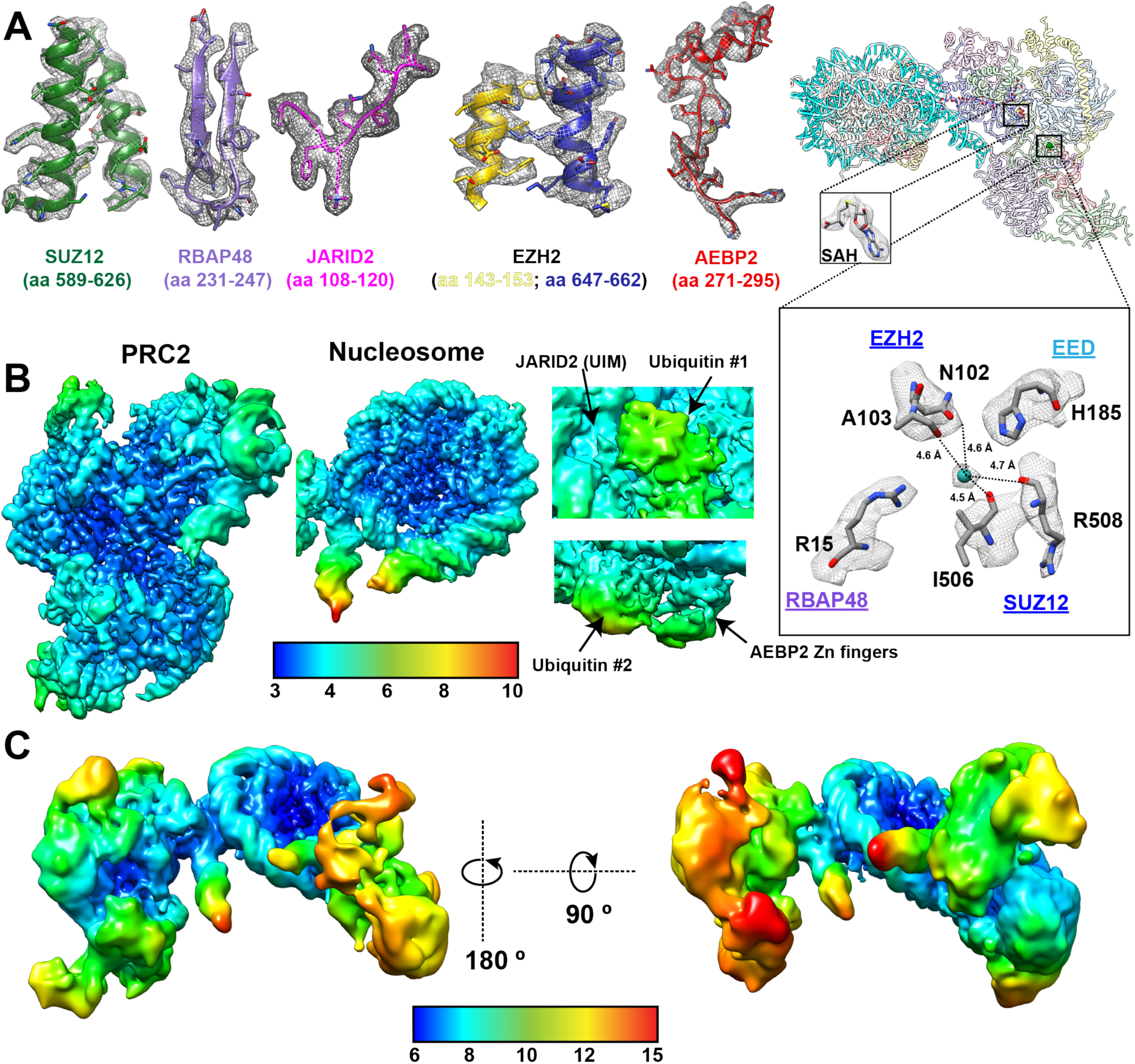
**(A)** Cryo-EM density and built in models for regions of the final map for PRC2-AEBP2-JARID2 (aa 1-450) bound to H2AK119ub containing mono-nucleosomes corresponding to the different proteins in PRC2. On the far right is the full model of PRC2-nucleosome with a close up view of two regions of PRC2, one showing the density for SAH and the corresponding atomic model of SAH and the other in the center of PRC2 showing density for a presumed divalent metal ion that appears to interact with regions from the four core subunits of PRC2 (EZH2, EED, SUZ12, and RBAP48). (B) Local resolution for PRC2 and for the nucleosome shown in high threshold and the close-up views of the local resolution of both ubiquitin molecules, JARID2 UIM and the AEBP2 tandem zinc fingers in lower threshold in the final reconstructions. (C) Local resolution of the map containing two PRC2 bound to a nucleosome.

**Fig. S4.**
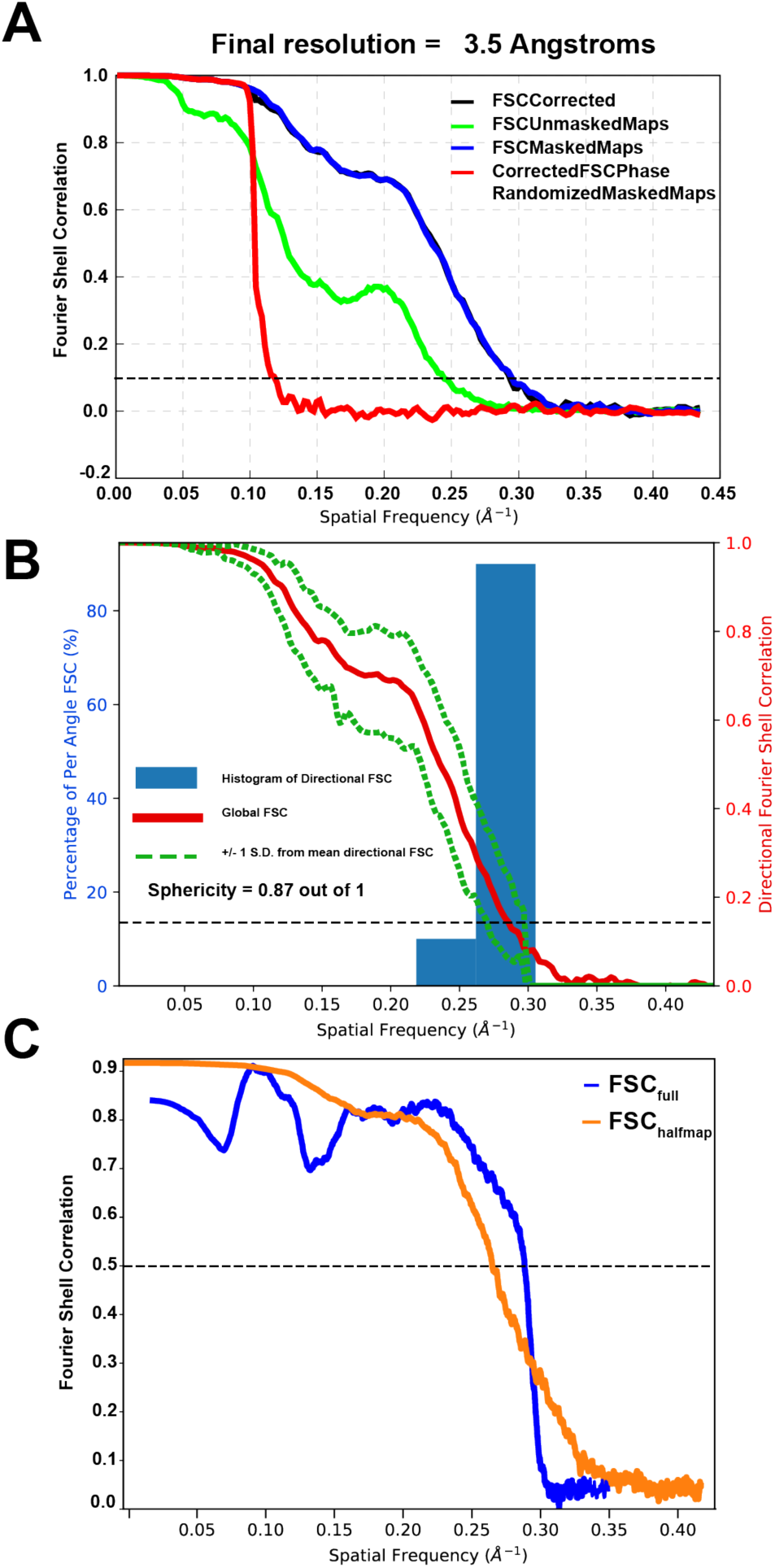
(A) FSC curves for the final, refined model. (B) 3D FSC for final Phenix-refined model vs full map. (C) Model vs Map FSC for final Phenix-refined model versus full map (Blue) and half-maps (orange). FSC/3D FSC.

**Fig. S5.**
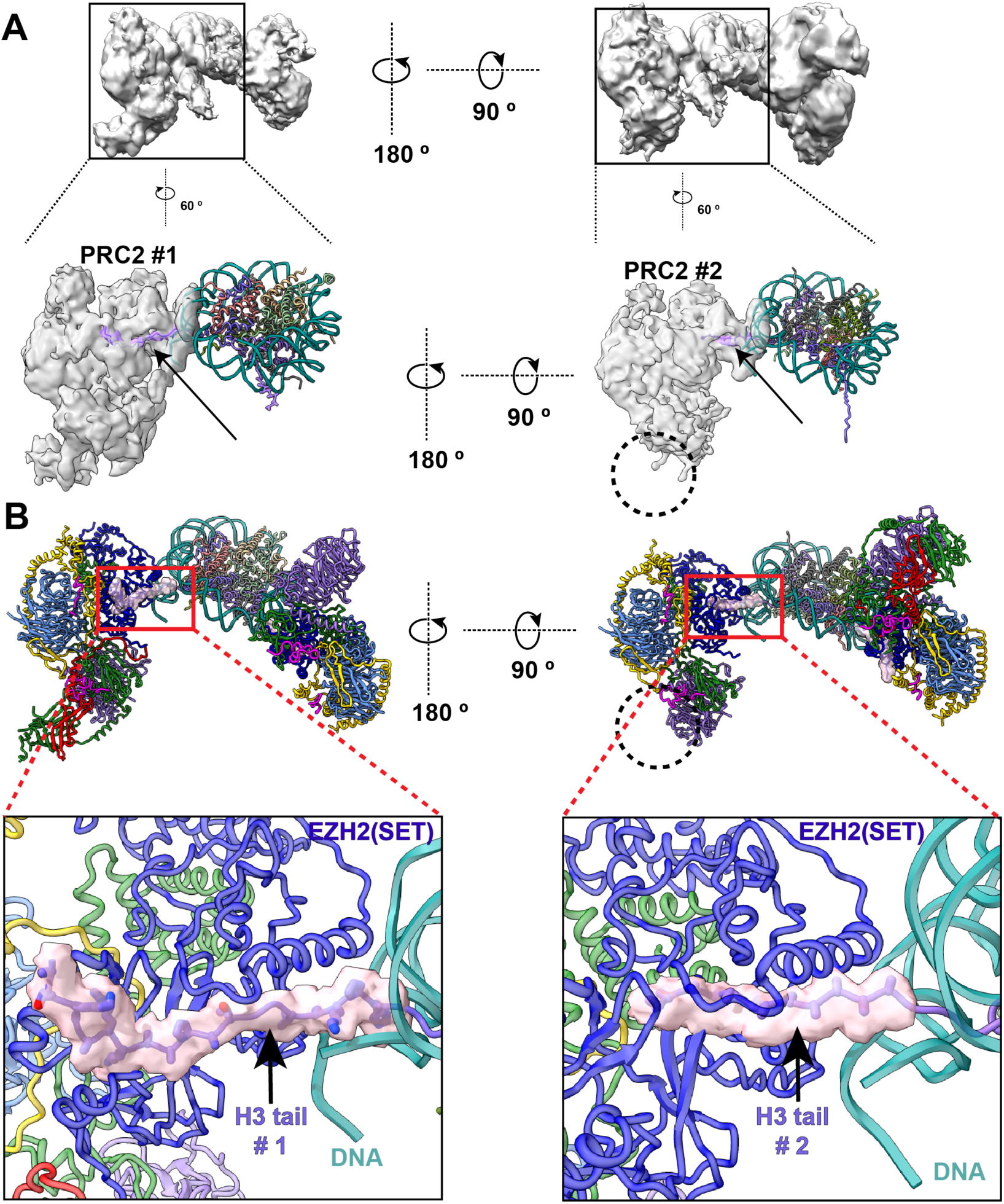
(A) (Top) The cryo-EM reconstruction of two PRC2-AJ_1-450_ engaging one each of the two histone H3 tails, on the opposite sides of the Ncl-Ub. (Bottom) The density map for each individual PRC2-ApJ_1-450_ complex, with the atomic models of the nucleosome and the H3 tails (pink) docked into the density. (B) (Top) Full model of the two PRC2-AJ_1-450_ engaging the nucleosome, with histone H3 density shown in pink. (Bottom) Close up view of the density for the histone H3 tails. Each PRC2 simultaneously interacts with one of the histone H3 tails (density shown in pink with model in stick representation).

**Fig. S6.**
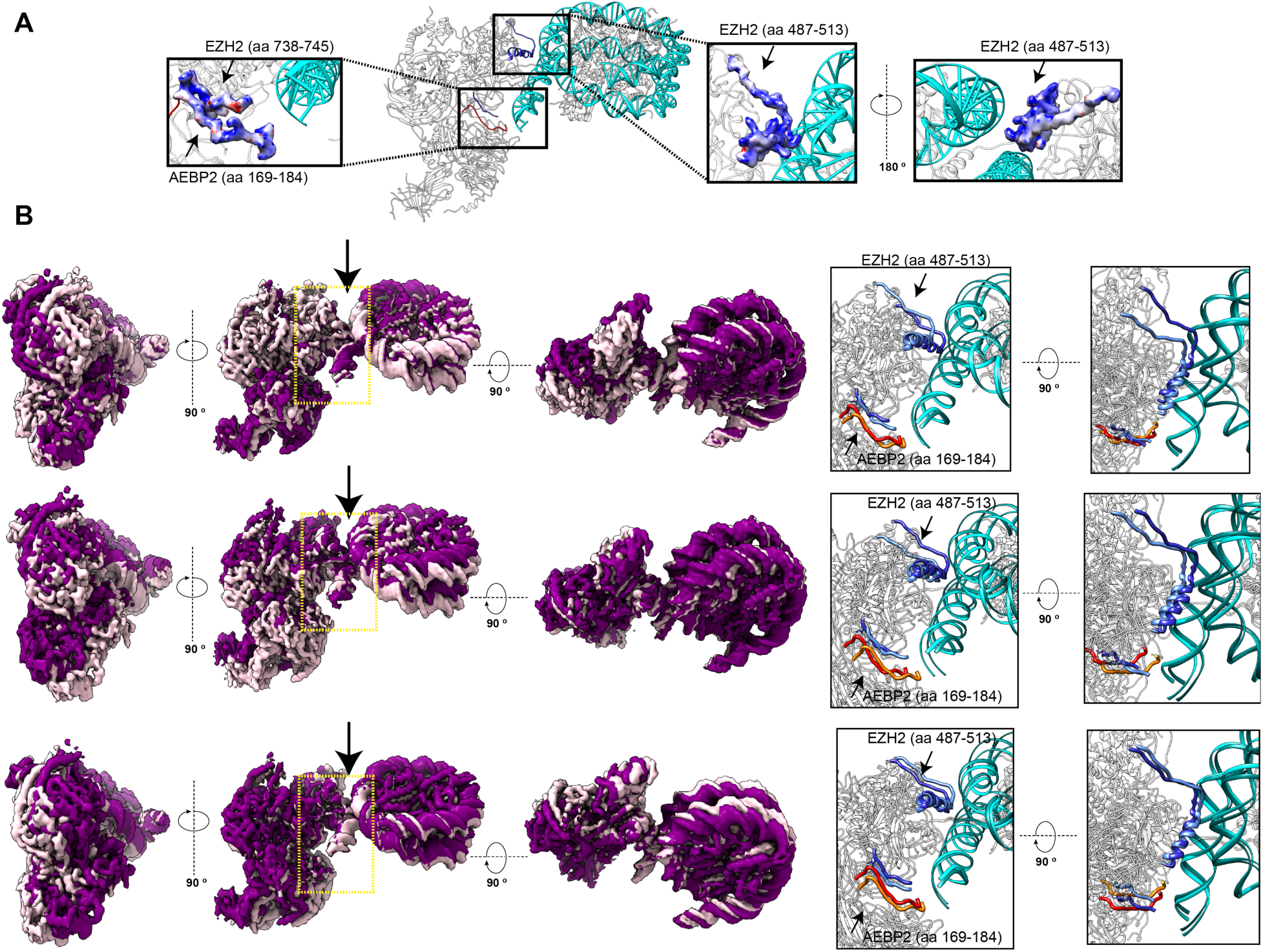
(A) Close-up view at the electrostatic potential of the EZH2 bridge helix region (aa 487-513) and an AEBP2 segment rich in lysine and arginine (aa 169-184) that interact with DNA showing the presence of strong positively charged surfaces in both segments. (B) Conformational flexibility observed using multibody analysis in RELION(*1*) in which PRC2 and nucleosome were treated as independent bodies. The three rows depict, in three orthogonal orientations, the representative conformational variance for the first three eigenvectors (63%). They show that the interface of PRC2-nucleosome interaction (black arrow) is flexible. On the far right, shown in two orthogonal orientations, are corresponding close ups of the nucleosome interacting regions (corresponding to the yellow dashed box on the left). The EZH2 bridge helix (aa 487-513) and C-terminus (aa 738-745) are shown in dark blue/light blue, the AEBP2 lysine and arginine rich segment (aa 169-174) in red/orange, and the nucleosomal DNA, shown in dark cyan/cyan. The dark-to-light shades of the respective colors illustrate the range of movement of these regions.

**Fig. S7.**
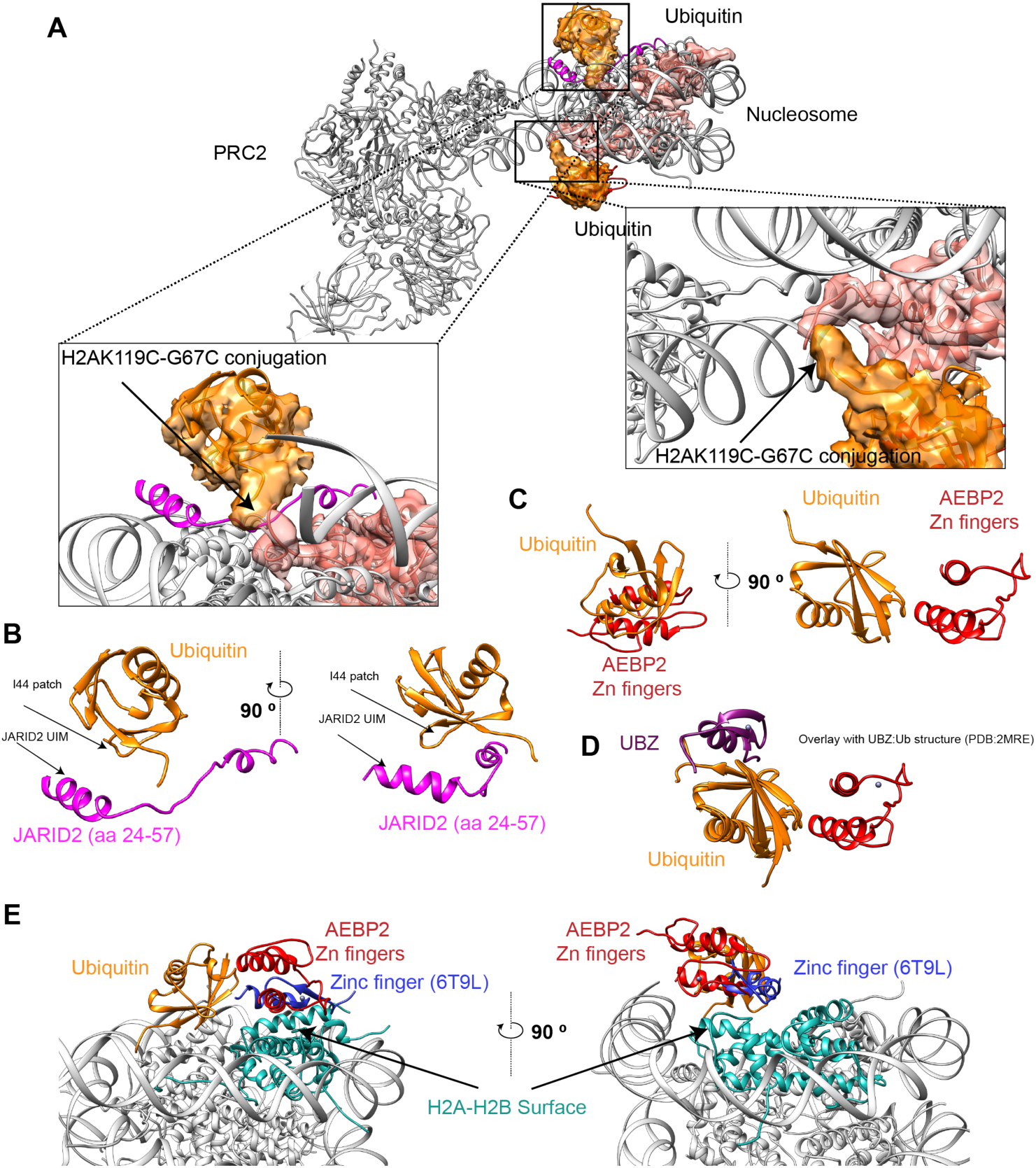
**(A)** Structure of PRC2-AJ_1-450_ bound to Ncl-ub showing the cryo-EM density for ubiquitin (orange) and H2A (salmon) (filtered based on local resolution). Close-up view showing the structure of ubiquitin rigid-body docked into the density such that the C-terminal tail of ubiquitin containing G76C is ∼ 3 Å from the H2AK119C. (B) Model of the JARID2 (aa 24-57) interaction with ubiquitin view in the same orientation as the close-up in (A), showing that I44-containing segment of ubiquitin is facing the JARID2 UIM. (C) Model of the ubiquitin-AEBP2 zinc finger interaction view in the same orientation as the close-up in (A) (left) and rotated 90 ° (right). (D) Superposition of the model of ubiquitin-AEBP2 zinc finger from this study with the NMR structure of the ubiquitin binding zinc finger (purple) bound to ubiquitin (orange; PDB:2MRE(*2*)) illustrating that the AEBP2 zinc finger does not interact with the canonical ubiquitin surface. (E) Superposition of the model of the ubiquitin-AEBP2 zinc finger from this study with the zinc finger (blue) from SAGA DUB module bound to ubiquitinated nucleosome (PDB:6T9L(*3*)) showing that they interact with the same H2A-H2B surface. The comparison was performed after rigid body docking the nucleosome in 6T9L within the cryo-EM density of PRC2-AJ_1-450_ bound to Ncl-ub. Only the zinc finger from 6T9L is shown for clarity.

**Fig. S8.**
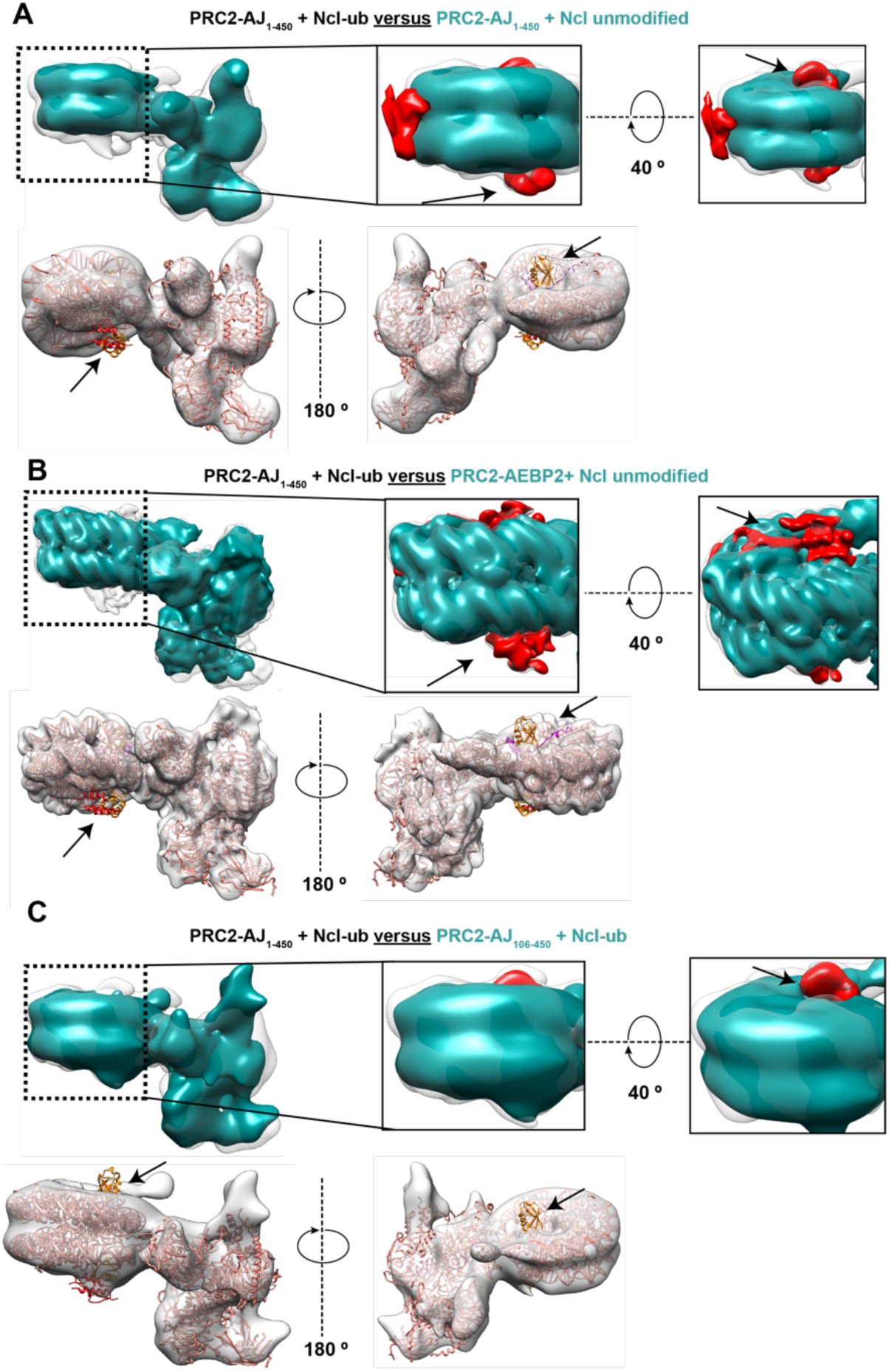
(A) (Top) Overlay of PRC2-AJ_1-450_ cryo-EM maps bound to a H2AK119ub-containing nucleosome (transparent map) and to an unmodified nucleosome (dark cyan). Close up views of the nucleosome with the difference map density for JARID2, AEBP2 zinc finger and ubiquitin are shown in red (indicated by black arrow). Both maps were lowpass filtered to 15 Å for comparison. (Bottom) Model of PRC2-AJ_1-450_ bound to H2AK119ub-containing nucleosome docked into the cryo-EM map of PRC2-AJ_1-450_ bound to unmodified nucleosome (transparent light grey) with arrows indicating ubiquitin, JARID2 and AEBP2 zinc finger for which no density is visible. (B) (Top) Overlay of PRC2-AJ_1-450_ cryo-EM maps bound to a H2AK119ub-containing unmodified nucleosome (transparent map) and PRC2-AEBP2 bound to an unmodified nucleosome (dark cyan). Close up views of the nucleosome with the difference map density for AEBP2 zinc finger and ubiquitin shown in red (indicated by the black arrow). Both maps were lowpass filtered to 8 Å for comparison. (Bottom) Model of PRC2-AJ_1-450_ bound to H2AK119ub-containing nucleosome docked into the cryo-EM map of PRC2-AEBP2 bound to an unmodified nucleosome (transparent light grey) with arrows indicating ubiquitin, JARID2 and AEBP2 zinc finger for which no density is visible. (C) (Top) Overlay of the cryo-EM reconstructions of PRC2-ApJ_106-450_ bound to H2AK119ub nucleosome (dark cyan) with that of PRC2-AJ_1-450_ bound to H2AK119ub containing nucleosome (transparent map). Close up views of the nucleosome with difference map density for the ubiquitin that binds JARID2 shown in red (indicated by black arrow). Both maps were filtered to 18 Å for comparison. (Bottom) Model of PRC2-AJ_1-450_ bound to H2AK119ub-containing nucleosome docked into the cryo-EM map of PRC2-AJ_106-450_ bound to a H2AK119ub unmodified nucleosome (transparent light grey) with arrows indicating ubiquitin and JARID2 for which no density is visible.

**Fig. S9.**
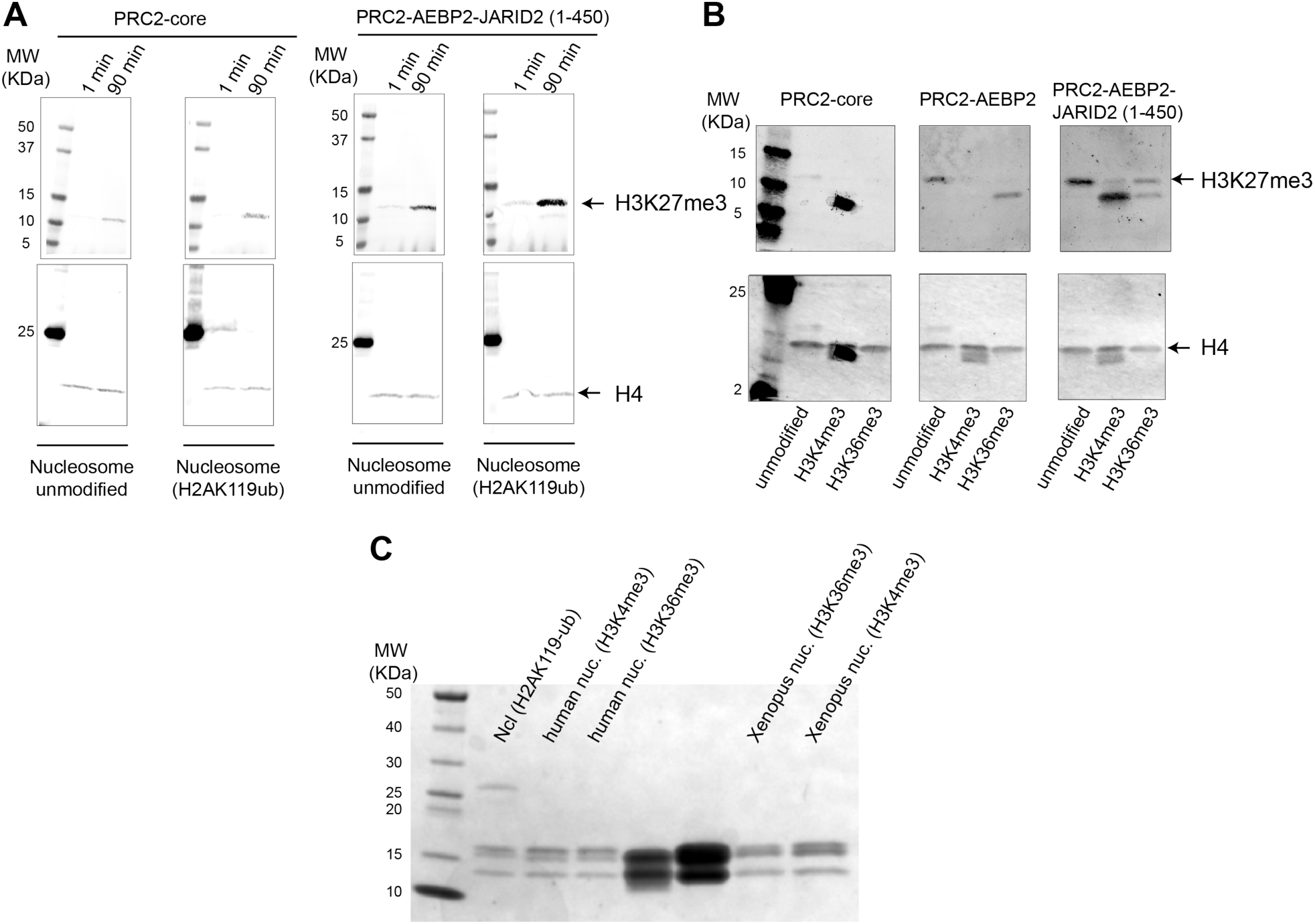
(A) Raw images of the western blot activity assay for PRC2-core and PRC2-AEBP2-JARID2 (aa 1-450) with either unmodified nucleosome or H2AK119ub containing nucleosome. Top panel shows Cy5 fluorescence images for H3K27me3 while bottom panel shows Cy3 fluorescence image for H4. (B) Raw images of the western blot activity assay for PRC2-core, PRC2-AEBP2, and PRC2-AEBP2-JARID2 (aa 1-450) with Xenopus nucleosomes unmodified, H3K4me3, and H3K36me3. Top panel shows Cy5 fluorescence images for H3K27me3 while bottom panel show Cy3 fluorescence images for H4. (C) Coomassie stained 4-12% Tris-Glycine SDS-PAGE gels of different nucleosomes used showing the presence of all four histones.

**Fig. S10.**
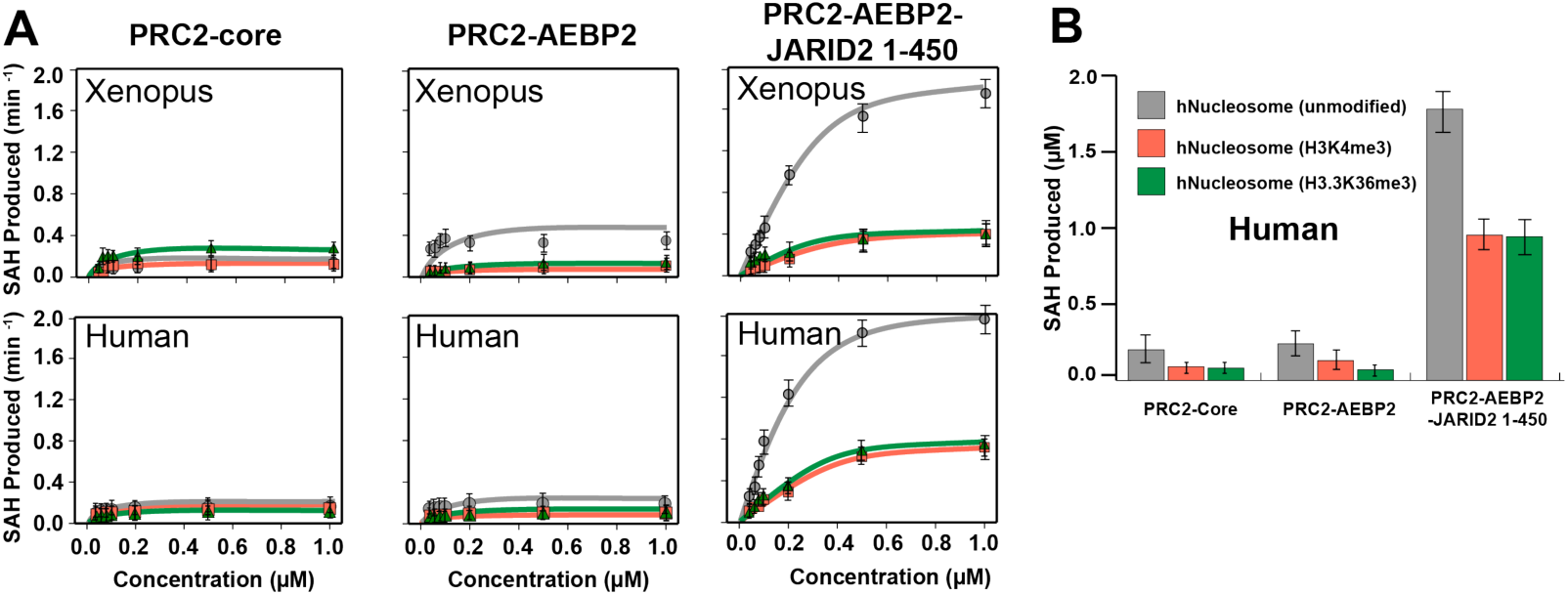
(A) Comparison of HMTase activity of PRC2 on Xenopus versus Human H3K4/K36me3 containing nucleosomes. Kinetic assays showing initial rates of four different PRC2 complexes with increasing amounts of either Xenopus nucleosomes either unmodified (grey) or containing H3K4me3 (orange) or H3K36me3 (green). (Bottom) As top row, but using Human nucleosomes. (B) Bar graph showing end point assays for cumulative H3K27me/me2/me3 activity for different PRC2 complexes on Human nucleosomes: unmodified (grey), H3K4me3-containing (orange), H3K36me3-containing (green).

**Fig. S11.**
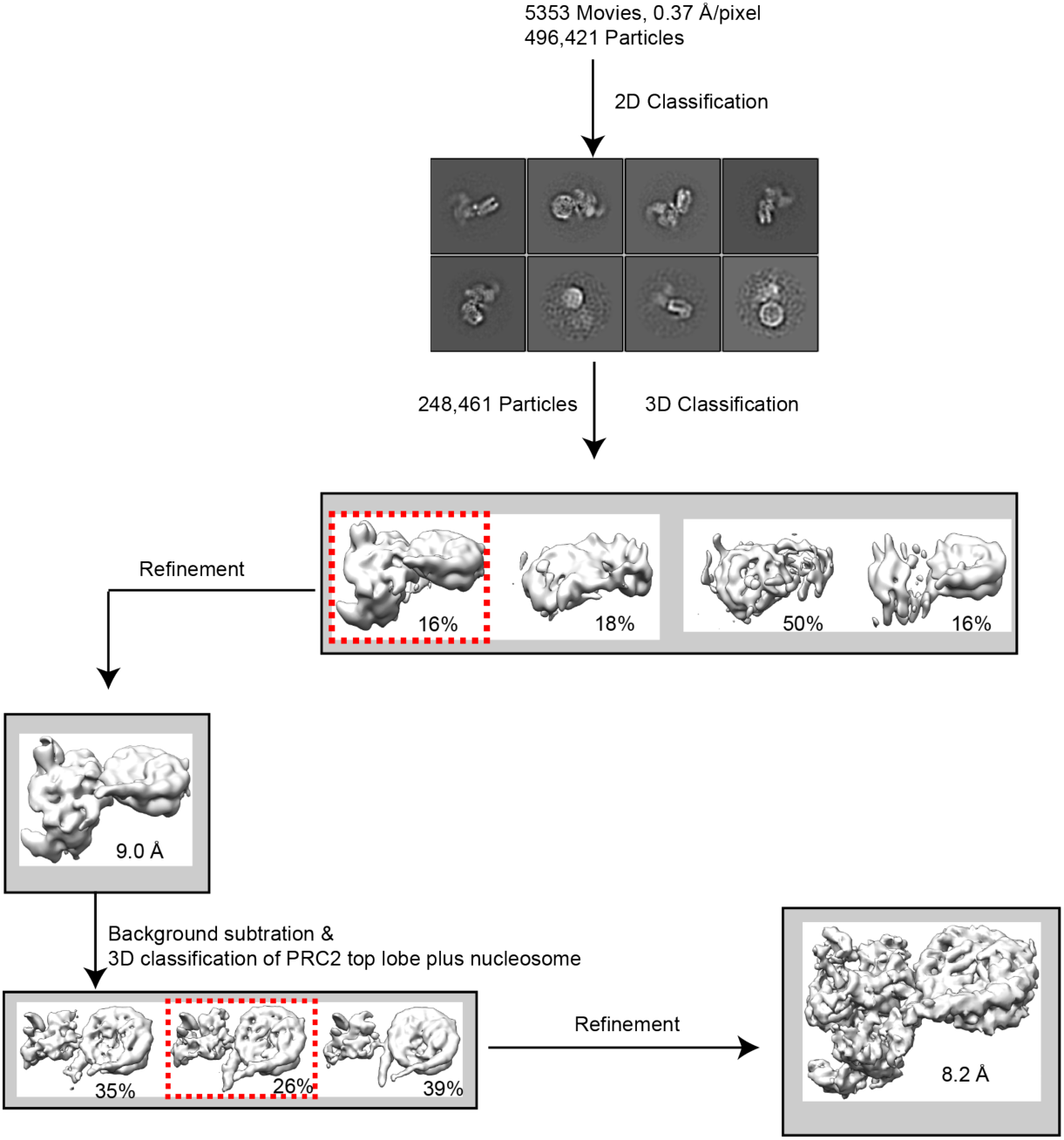
Processing work flow for PRC2-AEBP2-JARID2 (aa 1-450) bound to human H3K4me3-containing mono-nucleosome collected on a Titan Krios with Gatan K3 camera.

**Fig. S12.**
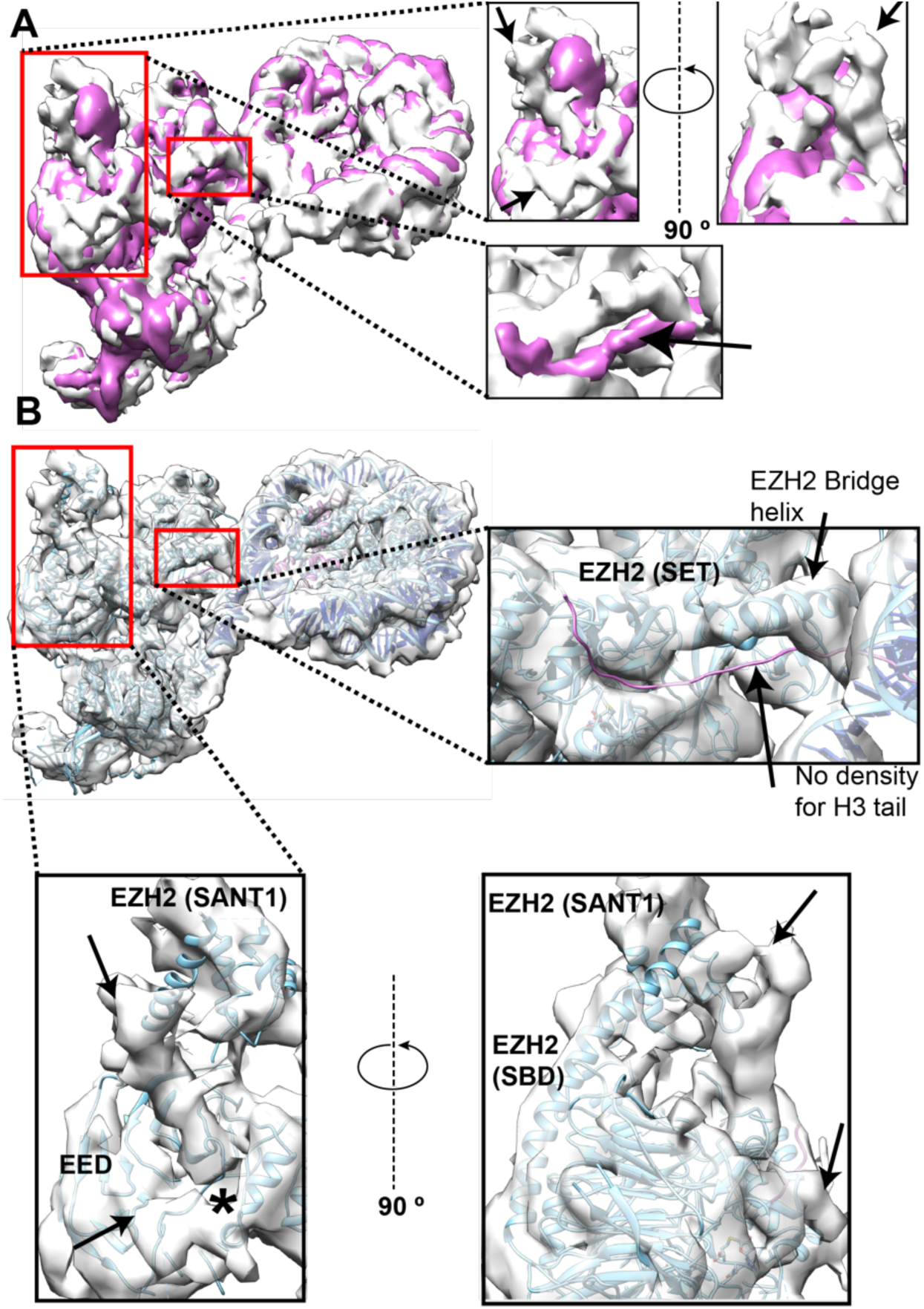
(A) Overlay of cryo-EM density maps for PRC2-AJ_1-450_ bound to human H3K4me3 (light grey) and to Xenopus Ncl-ub (purple) nucleosomes. Both maps are lowpass filtered to 8 Å for comparison. At the displayed threshold (=0.03) the density for ubiquitin, the AEBP2 zinc-fingers and the JARID2 UIM are not visible in the Ncl-ub map. The red boxes highlight regions of notable differences and are shown in close-up views on the right. (Top inset) Orthogonal close-up views of the EED, EZH2 (SANT1, SBD) region and JARID2 K116me3 binding region, with differences in the PRC2 bound to H3K4me3-containing nucleosome indicated by black arrows. (Bottom inset) Close-up view of the EZH2 (SET) catalytic domain showing that the density for H3 tail is observed in the PRC2-Ncl-ub reconstruction but it is absent in that for PRC2 bound to the H3K4me3-containing nucleosome. (B) The atomic model for PRC2 and nucleosome (cyan) from this study was fitted into the cryo-EM reconstruction of PRC2-AJ_1-450_ bound to Human H3K4me3 nucleosome (light grey) after removing ubiquitin, the JARID2 (UIM) and the AEBP2 zinc-fingers. The docked model shows that the overall geometry of interaction between PRC2 and the two types of nucleosome is the same. Flexible fitting was used in iMODFIT(*4*) to account for the differences in the EZH2 (SANT1, SBD) region. The SBD of EZH2 adopts a straight conformation and repositions the SANT1 domain, as also observed in one of the two active states in our earlier study of the PRC2-AEBP2-JARID2 structure without nucleosome, and in our previous structure of PRC2-AEBP2 bound to a dinucleosome(*5, 6*). (Top inset) Close-up view of the EZH2 (SET) domain showing the absence of density for the histone H3 tail in the catalytic region. (Bottom inset) Close-up view of the EED, EZH2 (SANT1, SBD) region and JARID2 K116me3 binding region showing unaccounted density (indicated by black arrow) observed in PRC2 bound to H3K4me3-containing nucleosome. The unaccounted density is near the region on EED where JARID2 K116me3 or H3K27me3 bind, but it occupies a larger volume and does not appear to overlap significantly with the JARID2 K116me3 segment (indicated by *) observed in the PRC2-AJ_1-450_ bound to Ncl-ub.

**Table S1.**
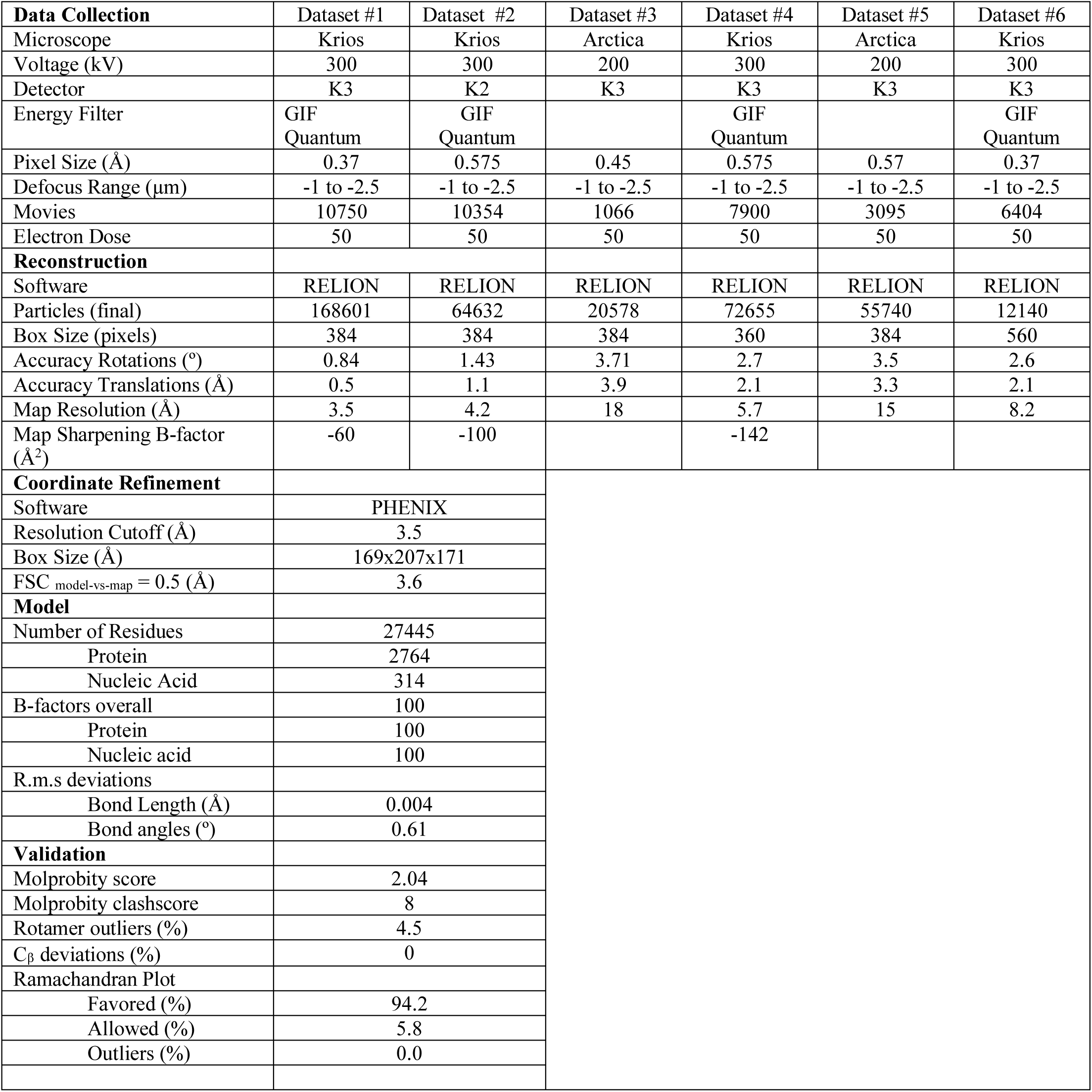
Cryo-EM data collection, refinement and validation statistics for the different data sets for the different data sets - #1: PRC2-AEBP2-JARID2 1-450+Ncl-ub, #2: PRC2-AEBP2-JARID2 1-450+Ncl-ub; #3-PRC2-AEBP2-JARID2 1-450+Ncl; #4:PRC2-AEBP2+Ncl; #5:PRC2-AEBP2-JARID2 106-450+Ncl-ub; #6: PRC2-AEBP2-JARID2 1-450+human Ncl (H3K4me3). Only the combined map from dataset #1 and #2 were used for model building.

**Movie 1** - Cryo-EM density map (3.5 Å overall resolution) of PRC2-AJ_1-450_ bound to Ncl-Ub, segmented with colors corresponding to the different proteins and/or domains of PRC2 and the nucleosome (as in Fig. 1), with the corresponding atomic model, shown in ribbon, for the whole assembly (same color scheme).

